# WACS: Improving ChIP-seq Peak Calling by Optimally Weighting Controls

**DOI:** 10.1101/582650

**Authors:** Aseel Awdeh, Marcel Turcotte, Theodore J. Perkins

## Abstract

**Motivation:** Chromatin immunoprecipitation followed by high throughput sequencing (ChIP-seq), initially introduced more than a decade ago, is widely used by the scientific community to detect protein/DNA binding and histone modifications across the genome. Every experiment is prone to noise and bias, and ChIP-seq experiments are no exception. To alleviate bias, the incorporation of control datasets in ChIP-seq analysis is an essential step. The controls are used to account for the background signal, while the remainder of the ChIP-seq signal captures true binding or histone modification. However, a recurrent issue is different types of bias in different ChIP-seq experiments. Depending on which controls are used, different aspects of ChIP-seq bias are better or worse accounted for, and peak calling can produce different results for the same ChIP-seq experiment. Consequently, generating “smart” controls, which model the non-signal effect for a specific ChIP-seq experiment, could enhance contrast and increase the reliability and reproducibility of the results.

**Results:** We propose a peak calling algorithm, Weighted Analysis of ChIP-seq (WACS), which is an extension of the well-known peak caller MACS2. There are two main steps in WACS: First, weights are estimated for each control using non-negative least squares regression. The goal is to customize controls to model the noise distribution for each ChIP-seq experiment. This is then followed by peak calling. We demonstrate that WACS significantly outperforms MACS2 and AIControl, another recent algorithm for generating smart controls, in the detection of enriched regions along the genome, in terms of motif enrichment and reproducibility analyses.

**Conclusion:** This ultimately improves our understanding of ChIP-seq controls and their biases, and shows that WACS results in a better approximation of the noise distribution in controls.

## Background

High throughput sequencing technologies help in uncovering the mechanisms of gene regulation and cell adaptation to external and internal environments [1, 2]. One widely used technology is chromatin immunoprecipitation followed by next generation sequencing (ChIP-seq). It allows the genome-wide investigation of the structural and functional elements encoded in a genomic sequence, such as transcriptional regulatory elements. The main goal of a ChIP-seq experiment is the detection of protein-DNA binding sites and histone modifications genome-wide in various cell lines and tissues. Many peak calling methods have been proposed for the identification of regions of enrichment (putative binding sites) in ChIP-seq data [3, 4, 5, 6, 7].

Every experiment is prone to noise and bias, and ChIP-seq experiments are no exception. While some read pileups correspond to regions of true enrichment, others may be a result of the distortion of the ChIP-seq signal. Biased or noisy datasets (with a high number of false negative or false positive peaks) negatively impact downstream biological and computational analyses [8]. Thus, accounting for both noise and bias is important. Existing peak callers generally account for noise by assessing statistical significance under some statistical model. Bias is a more complicated subject and is usually addressed explicitly only via some control data to which the ChIP-seq is compared. We return to the issue of controls shortly.

There are many sources of bias in a ChIP-seq experiment. In the experimental design, for example, the quality of the experiment is predetermined by anti-body and immunoprecipitation specificity. Low sensitivity, resulting from poor affinity to the target protein of interest, or low specificity, from cross reactivity with other unrelated proteins, degrades the quality of a ChIP-seq experiment [9]. The fragmentation step may also introduce bias [10]. Prior to immunoprecipitation, the DNA-protein complexes undergo fragmentation. However, due to the non-uniform nature of the chromatin structure (DNA), some regions are more densely packed (heterochromatin) than others and are thus more resistant to fragmentation. Less densely packed regions (euchromatin) will undergo more fragmentation. Another source of bias is mappability, which is the extent to which reads are uniquely mapped to regions along the genome [10, 11]. In an ideal situation, long enough reads are used such that there is higher coverage and uniformity in coverage. However, in practice, read length is short and there are “ambiguous” reads that map to multiple regions. Such reads can either be multiply mapped (creating false positive ChIP-seq signal) or discarded (creating empty, unmappable regions), with either choice creating a different sort of bias. GC content bias [12, 13], introduced by PCR amplification or sequencing, also results in imbalanced coverage of reads along the genome. For example, in PCR amplification, GC rich fragments are targeted more than the GC poor fragments. These variations in coverage can have a significant impact on the results obtained.

Systematic and experimental biases hinder the full potential of ChIP-seq analysis. Thus, the quality of the input samples is important, especially in large scale analysis where low quality datasets have greater effects [8, 14]. Consequently, more than a decade after ChIP-seq was introduced, the ENCODE and modENCODE consortia developed a set of ChIP-seq quality control metrics and guidelines to produce high quality reproducible data [9]. The protocols address all the stages of a ChIP-seq experiment, as bias and noise may be introduced at various stages, such as experimental design, execution, evaluation and storage methods [10].

One essential step for the alleviation of bias is the incorporation of control datasets in ChIP-seq analysis. It assists in the selection of true enrichment binding sites from false positives. Controls, such as input DNA and IgG, attempt to minimize the effects of immunoprecipitation, antibody imprecision, PCR-amplification, mappability bias, etc., and thereby increase the reliability of the results. In the input DNA, using the same conditions as the original ChIP-seq experiment, the DNA undergoes cross linkage and fragmentation. However, no antibody nor immunoprecipitation is used [9]. For the IgG control, sometimes referred to as a “mock” ChIP-seq experiment, all the same steps and conditions as the original ChIP-seq experiment are applied. However, a control antibody (not specific to the protein of interest) is adopted to interact with nonrelevant genomic positions [9]. DNase-seq and ATAC-seq are used to tackle open chromatin regions. According to ENCODE [9], the input DNA and IgG controls should have a sequencing depth greater than or equal to the original ChIP-seq experiment. Higher sequencing depth is recommended since input DNA signals represent broader genomic chromatin regions than ChIP-seq [9, 10]. Other crucial factors addressed by the protocols include, but are not limited to, biological/technical replicates and library complexity.

Many existing peak calling algorithms allow testing enrichment compared to a control [7, 15, 16, 17, 18, 19, 20]. Whether biases in controls and ChIP-seq data are the same is not known, however. None of these methods selects a control or estimates background signals. Depending on which controls are selected and their nature, peak callers can produce different results (i.e., binding site positions) for the same ChIP-seq experiment. The BIDCHIPS [21], CloudControl [22] and AIControl [23] studies have shown that different ChIP-seq datasets can be biased in different ways. They address different biases in different ChIP-seq datasets via the integration of multiple control datasets through regression to improve enrichment analysis. There are some limitations to these studies, however.

For example, BIDCHIPS [21] has the ability to reprioritize peaks already identified by another peak calling method. However, only five notions of control are accounted for and there are no mechanisms for de novo peak calling based on the combined control [21]. The Hiranuma et al. [22, 23] studies prove the advantage of using more controls to model the background signal. In CloudControl [22], the controls are subsampled in their regression fit proportional to their weights. This then allows the single customized control to be used as input to any peak calling method. However, the down-sampling of the combined controls may introduce noise into the control signal.

AIControl [23], a peak calling framework, is an extension of CloudControl [22]. It integrates a group of publicly available control datasets and uses ridge regression to model the background signal. This eliminates the need for the user to input controls. However, some users may want to provide their own controls, and this is not accommodated. Additionally, the number of datasets in ENCODE increases with time, so allowing controls as input in a weighted peak caller is important to represent the newly available datasets and newly explored cell lines.

In this work, we introduce a peak calling algorithm, Weighted Analysis of ChIP-Seq (WACS), which utilizes “smart” controls to model the non-signal effect for a specific ChIP-seq experiment. WACS first estimates the weights for each input control, without requiring the fine-tuning of any parameters. Using the weighted controls, WACS then proceeds to detect regions of enrichment along the genome. WACS is an extension of MACS2.1.1 (Model-based Analysis for ChIP-Seq) [18], the most highly cited open source peak caller. Our development of WACS based on MACS2 allows researchers to use the weighted approach within a peak calling method with which they are familiar, and which has many refined features. Fragment length estimation/detection, read shifting, candidate peak identification, and peak assessment remain the same, while the construction of the control via the weighted combination of datasets is different. To allow for potentially large numbers of controls, we restructure the code invisibly for better memory footprint. We also correct a hashing bug in the pileup-computing code of MACS2, which becomes especially important when we have high read depth and/or many controls. (This bug has subsequently been corrected in the main MACS2 distribution as well.)

We evaluate WACS on a large collection of 438 ChIP-seq datasets and 147 control datasets from the K562 cell line in the ENCODE database [24]. To establish generalizability and study performance in a less expansive setting, we also investigate WACS on 20 ChIP-seq datasets for each of the A549, GM12878 and HepG2 cell lines. (The terms ChIP-seq and treatment are used interchangeably throughout the paper.) The results demonstrate the importance of smart bias removal methods and the use of customized control datasets for each ChIP-seq experiment, as the amount of bias varies across different ChIP-seq experiments. In the investigation of downstream genomic analysis, such as motif enrichment and reproducibility, the use of weighted controls in WACS shows a significant improvement in peak detection in comparison with the pooled unweighted controls in MACS2 and weighted controls in AIControl.

### Algorithm 1 Derive Weights

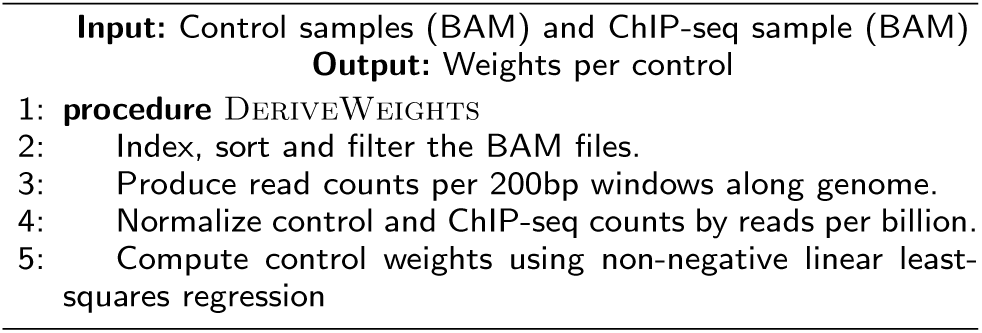

## Methods

### WACS: Weighted Analysis of ChIP-seq

Our approach, WACS, estimates a background distribution by weighting controls, and ultimately identifies regions of enrichment along the genome (Figure 1 and Supplementary Figure 1). Below we describe the five major steps of the WACS algorithm. To implement WACS, we modified a well-known open source algorithm, MACS2. Because there is limited written description of how MACS2 works, we describe some parts of MACS2 to fully describe WACS. The WACS algorithm is summarized into two parts: Derive Weights (Algorithm 1) and Peak Detection (Algorithm 2).

**Figure 1:**
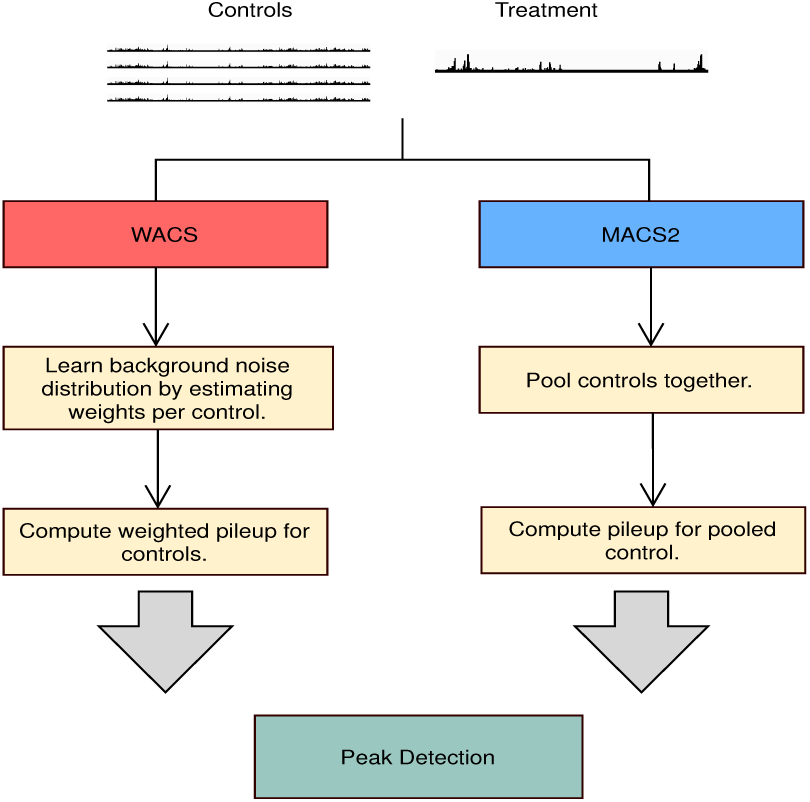
Flowcharts for WACS and MACS2. Both methods take controls and a treatment as input.

### Algorithm 1: Derive Weights

The control and treatment samples (in BAM format) are first preprocessed, as seen in Algorithm 1. Using SAMtools [25], we index, sort and optionally filter (remove duplicates from) the BAM files (line 2 in Algorithm 1). We then use BEDtools [26] to convert the BAM files of mapped reads into read counts per 200 base pair (bp) windows along the genome with 50 bp increments (line 3 in Algorithm 1).

Next, WACS normalizes the mapped reads per window for the preprocessed control and treatment samples. This ensures that the control and treatment samples are on the same scale. WACS applies reads per billion normalization to both the control and ChIP-seq samples (line 4 in Algorithm 1). For each sample *m* and window *i*:

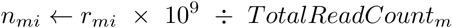

where *r*_*mi*_ is the read count in the window, *n*_*mi*_ is the normalized read count, and *TotalReadCount*_*m*_ is the total number of reads in sample *m*. This effectively reproduces the normalization in MACS2, which linearly scales the control sample to the ChIP-seq sample. In what follows, we assume *k* total controls comprise samples 1 to *k*, and sample *k* + 1 is the ChIP-seq data.

#### Algorithm 2 Peak Detection

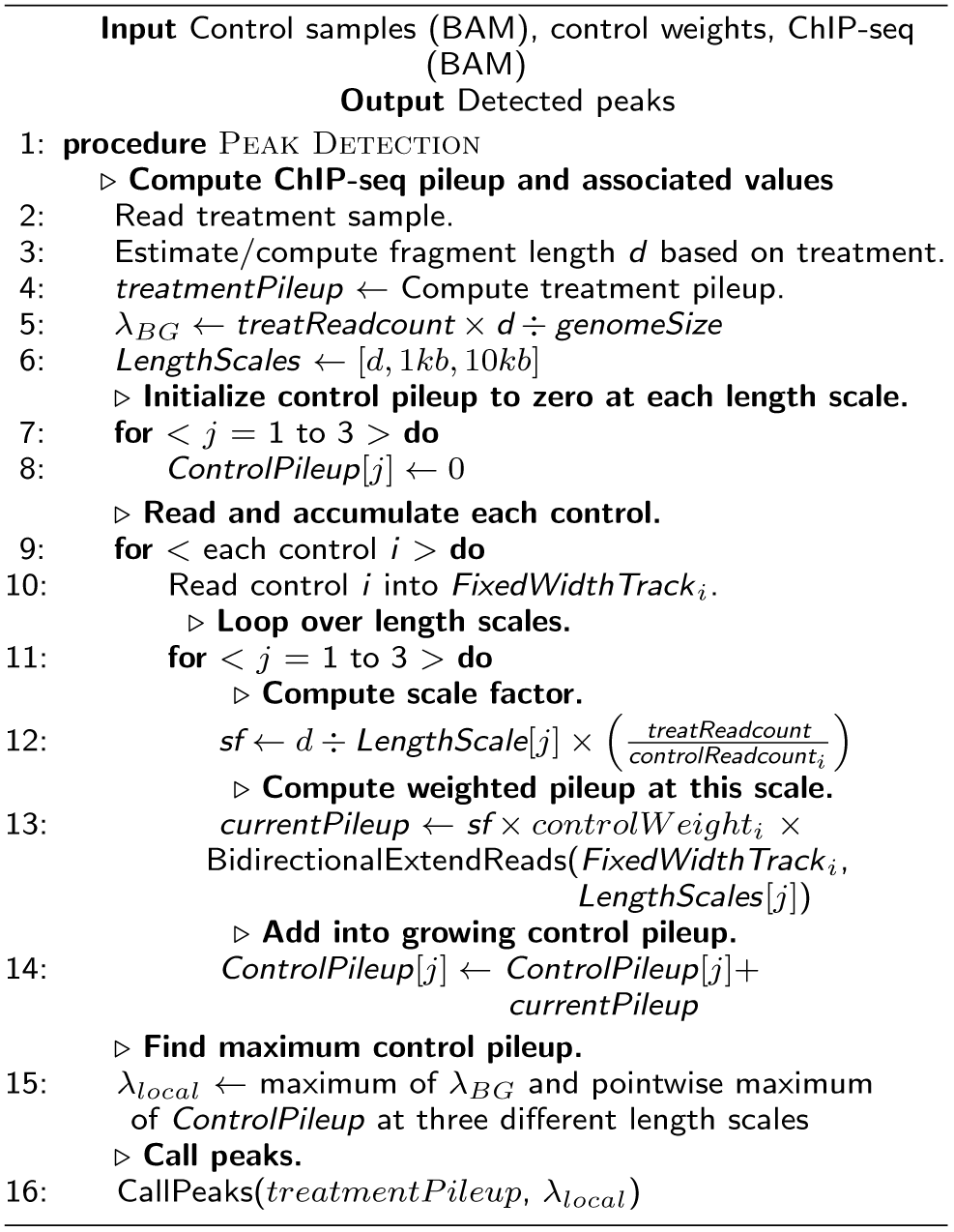

WACS then calculates the weights per input control (line 6 in Algorithm 1). WACS performs non-negative least squares (NNLS) to model the treatment dataset as a function of the controls. The overall objective of the regression is to find the values of the parameters (weights), that minimize the sum of squared differences between predictions and target values, with an additional constraint that allows only positive weights. Given *n* instances (windows), *y*_*i*_ = *n*_*k*+1,*i*_ target values (one per window), *x*_*i*_ = (*n*_1*i*_, *…, n*_*ki*_) feature vectors (one vector per window), a vector Θ of coefficient weights and a constant offset Θ_0_, NNLS’s objective function is:

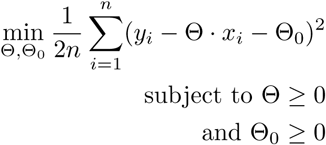

To solve the NNLS regression we rely on the nnls module from scipy.optimize, part of the scipy [27] package in Python. This produces a weighted control model for the treatment, with weights that indicate the relative importance of each control in modelling the treatment background signal. Zero weights are given to controls not required for modelling the treatment experiment. If there is one control, WACS and MACS2 produce the same output, as by default, the control in WACS gets a weight of exactly 1. The controls can also be weighted by the user, instead of using NNLS to compute the weights of the controls.

### Algorithm 2: Peak Detection

WACS is identical to MACS2 in its initial processing of the treatment sample, including: loading the mapped reads (line 2); estimation/calculation of fragment length *d*, which differs depending on whether the ChIP-seq reads are sequenced single-end or paired-end (line 3); and construction of the treatment pileup, which also differs for single-end or paired-end reads (line 4). Because these details have been described elsewhere, we do not repeat them here [18, 28, 29].

Where WACS differs substantially from MACS2 is how it reads in, processes, and combines the control samples. WACS reads the controls into memory one at a time, accumulating them into overall (weighted) control pileups at three different length scales: *d*, 1*kb* and 10*kb*. The length scale is essentially the diameter of a Parzen-windows density estimator used to smooth the control reads. As each control is read in, it is smoothed, scaled so that its total reads are commensurate with the treatment, and further scaled by the control weight computed in Algorithm 1 (unless the user opts for unweighted controls). The function BidirectionExtendReads performs the actual smoothing, extending the read starts into intervals with diameter equal to the length scale. The smoothed and scaled control is added to the growing overall control at that length scale. In contrast, MACS2 reads all the control data in before beginning smoothing, which can create an unmanageable memory footprint when very many controls are being combined. Finally, WACS (as does MACS) creates an overall control pileup by taking the pointwise maximum of the “background” read density *λ*_*BG*_ and the control pileups computed at each length scale.

Finally, WACS calls peaks using the same mechanism as MACS2, which involves identifying candidate peaks and comparing the pileup heights at their summits with the control track. In the case of unweighted controls, WACS produces an identical control track to MACS2 and identical peak calls. However, when control samples are weighted differently, a different control track is produced and different peaks may be called. Each peak is associated with a p-value and a q-value, the latter accounting for multiple comparisons across the entire genome.

### Duplicate removal

Duplicate reads—multiple reads mapped to the same position on the genome—are often due to the overamplification of DNA fragments by PCR, which leads to the repeated sequencing of a DNA fragment. For WACS and MACS2, duplicate removal is optional. To produce more reliable peak calls, MACS2/WACS remove redundant reads at each genomic locus for both the treatment and control datasets [18].The default number per genomic locus is determined by the sequencing depth. However, when dealing with multiple controls, MACS2 performs duplicate removal after pooling reads. WACS does the same thing when used in unweighted mode, for the sake of consistency with MACS2. In this case, apparent “duplicates” arising from different sequencing runs may be removed incorrectly, artificially flattening the control read distribution in high density areas. This phenomenon can be particularly prominent when hundreds of controls are being pooled. Thus, we recommend that users who want to perform de-duplication do so prior to feeding the mapped read files to MACS2 or WACS.

### Experimental Evaluation

We evaluated WACS, MACS2.1.1 (https://github.com/taoliu/MACS) and AIControl (https://github.com/hiranumn/AIControl.jl/) on 438 ChIP-seq (treatment) and 147 control samples from the K562 cell line; and 20 ChIP-seq and 20 control samples for each of A549, GM12878, and HepG2 cell lines. (See Supplementary Tables 1, 2, 4, 5, and 6 for the accession codes of samples). As seen in Figure 1 (and Supplementary Figure 1), MACS2 pools the controls together for each ChIP-seq sample, whereas WACS estimates a weight for each control and computes a unique weighted control pileup for each ChIP-seq sample. AIControl uses a predefined set of publicly available controls [23]. For each ChIP-seq sample, we generated peaks under four conditions: (1) MACS2 with all the controls from the same cell line (All MACS2), (2) MACS2 with the matched ENCODE controls (Matched MACS2), (3) WACS with all the controls from the same cell line (WACS) and (4) AIControl with its predefined controls (AIControl).

Next, we used two methods to compare of the quality of the peaks generated by WACS, MACS2 and AIControl. One method considers all the original peaks output by each algorithm (called All Peaks). However, different peak callers can produce peaks in different locations based on the same data, and they can also produce different numbers of peaks. Thus, for additional comparison, we adopted the standardization procedure proposed by Hiranuma et al. [23], where the peak width and number of peaks are normalized for each treatment sample. First, the peak width is normalized by binning the peaks in 1000 base pair windows. For example, a peak at chromosome 1 from 14520 to 15420 is counted as two peaks covering bins 14000 to 15000 and 15000 to 16000. Next, the number of peaks under all four conditions for the same dataset is normalized. The top *n* most statistically significant peaks are selected for each of the peak-width normalized datasets, such that *n* is the minimum number of peaks under the four conditions.

## Results

### Peaks identified by WACS are more enriched for known sequence motifs

The purpose of ChIP-seq analysis is the identification of regions of enrichment, such as TF binding sites, along the genome. To evaluate the performance of our method in comparison to MACS2 and AIControl, we performed motif enrichment analysis. In this and the following two subsections, we focus on the K562 results, with results in additional cell lines reported further below. Adopting a similar motif analysis method as in [23], we first used JASPAR to obtain position weight matrices (PWMs) for each unique TF [30]. (See Supplementary Table 3 for the PWM IDs per TF.) Using PWMs as input, we then used FIMO (Find Individual Motif Occurrences) [31] in the MEME suite [32] to scan the genome and identify motif hits genome wide. We found motifs for 125 treatment samples. Motif hits tend to be enriched in genuine binding sites. In our analysis, peaks with a motif are considered as true positives, while those lacking a motif hit are considered false positives.

Fig 2a and 2b display the motif enrichment for each of these ChIP-seq datasets for all peaks and standardized peaks respectively, when using WACS (blue line), All MACS2 (red line), Matched MACS2 (green line) and AIControl (purple line). Using all the peaks, WACS outperforms All MACS2, Matched MACS2 and AIControl on 107 treatment samples in total. WACS also outperforms All MACS2, Matched MACS2 and AIControl on the majority of the treatment samples, when peaks are standardized, as shown in Table 1. Using a paired sign test with a two-tailed null hypothesis to compare the precision of WACS with MACS2 and AIControl, we get a p-value of less than 10^−5^. The two versions of MACS2 (with all or matched controls) perform similarly to each other. Matched MACS2 slightly outperforms All MACS2, suggesting that the ENCODE selected controls are well-matched to the treatment sample. Across the treatment samples, AIControl has the highest fluctuations in the precision values. The four methods perform similarly on treatment samples when the precision is less than 0.1. This suggests that these treatment samples may need to be more thoroughly investigated. Possibly the sample are of poor quality, or the motifs in JASPAR are not representative of true binding preferences.

**Table 1:** Motif Enrichment Summary.

| Type | WACS | All MACS2 | Matched MACS2 | AIControl |
| --- | --- | --- | --- | --- |
| All Peaks | 107 | 8 | 10 | 0 |
| Standardized | 90 | 8 | 16 | 11 |

**Figure 2:**
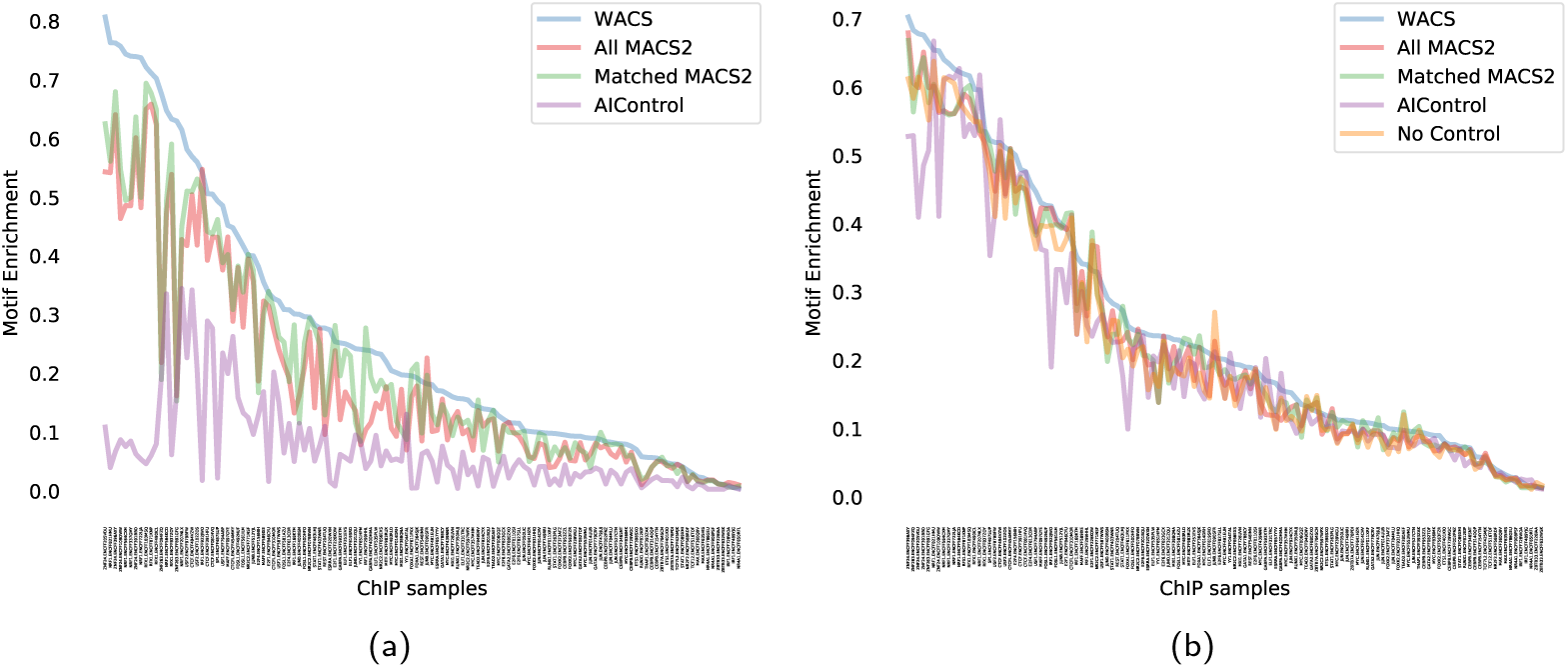
(a) Motif enrichment of the treatment samples, for each of the four peak calling methods. Motif enrichment (precision) is defined as the fraction of all peaks that contain at least one motif occurrence for the transcription factor in question. (b) Motif enrichment for the standardized peaks.

Another method for evaluating motif enrichment is the area under the precision-recall curve (AUPRC) [23]. The AURPC is designed to compare algorithms on the same set of instances. Each algorithm, however, generates a different set of peaks for a specific ChIP-seq dataset. Thus, we believe precision is a more appropriate evaluation metric than AUPRC for this comparison. Nevertheless, for the purpose of comparison with AIControl [23] which uses the AUPRC metric, we performed the AUPRC analysis as well. Supplementary Figure 2 shows an example precision-recall curve for the ChIP-seq dataset ENCFF109OWW with TF ZNF24, and Supplementary Figure 3 shows the the AUPRC for each of these ChIP-seq datasets when using standardized peaks. Using AUPRC, WACS outperforms All MACS2, Matched MACS2 and AIControl on 114, 99 and 100 of the 125 treatment samples respectively. These differences are statistically significant by a two-tailed sign test with p-value less than 10^−5^.

### Peaks identified by WACS are more reproducible

Ideally, a ChIP-seq peak calling algorithm is able to reproducibly identify true regions of enrichment along the genome with no false positives. Reproducibility is most commonly measured by computing the percentage overlap of peaks between replicates [4, 5]. According to the ENCODE guidelines, each ChIP-seq experiment is associated with at least two ChIP-seq biological replicate samples [24]. Here, we computed the percentage overlap between the non-standardized and standardized peaks separately for WACS, All MACS2, Matched MACS2 and AIControl across 197 ChIP-seq experiments with exactly two replicates. (The percentage overlap is equal to the overlap between the replicates divided by the total number of peaks from both replicates.) (See Table 2 in Supplementary Data for details.)

**Table 2:**
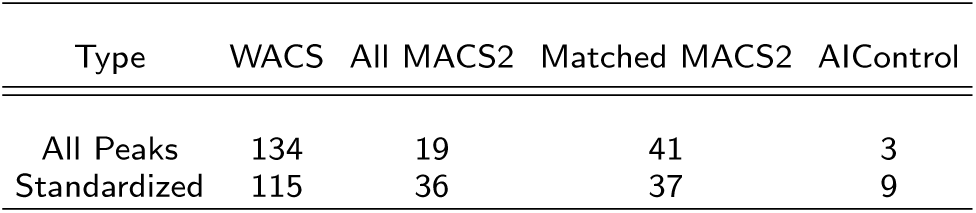
Reproducibility Analysis Summary.

Figure 3c and 3d show the percentage overlap with all peaks and standardized peaks respectively for each of the ChIP-seq experiments when using WACS (blue line), All MACS2 (red line), Matched MACS2 (green line) and AIControl (purple line). Using all peaks, WACS has higher reproducibility than All MACS2, Matched MACS2 and AIControl on 134 of the 197 ChIP-seq experiments. WACS also outperforms All MACS2, Matched MACS2 and AIControl on the majority of the ChIP-seq experiments with standardized peaks (Table 2). These differences are statistically significant using a paired sign test and at a p-value of less than 10^−5^.

**Figure 3:**
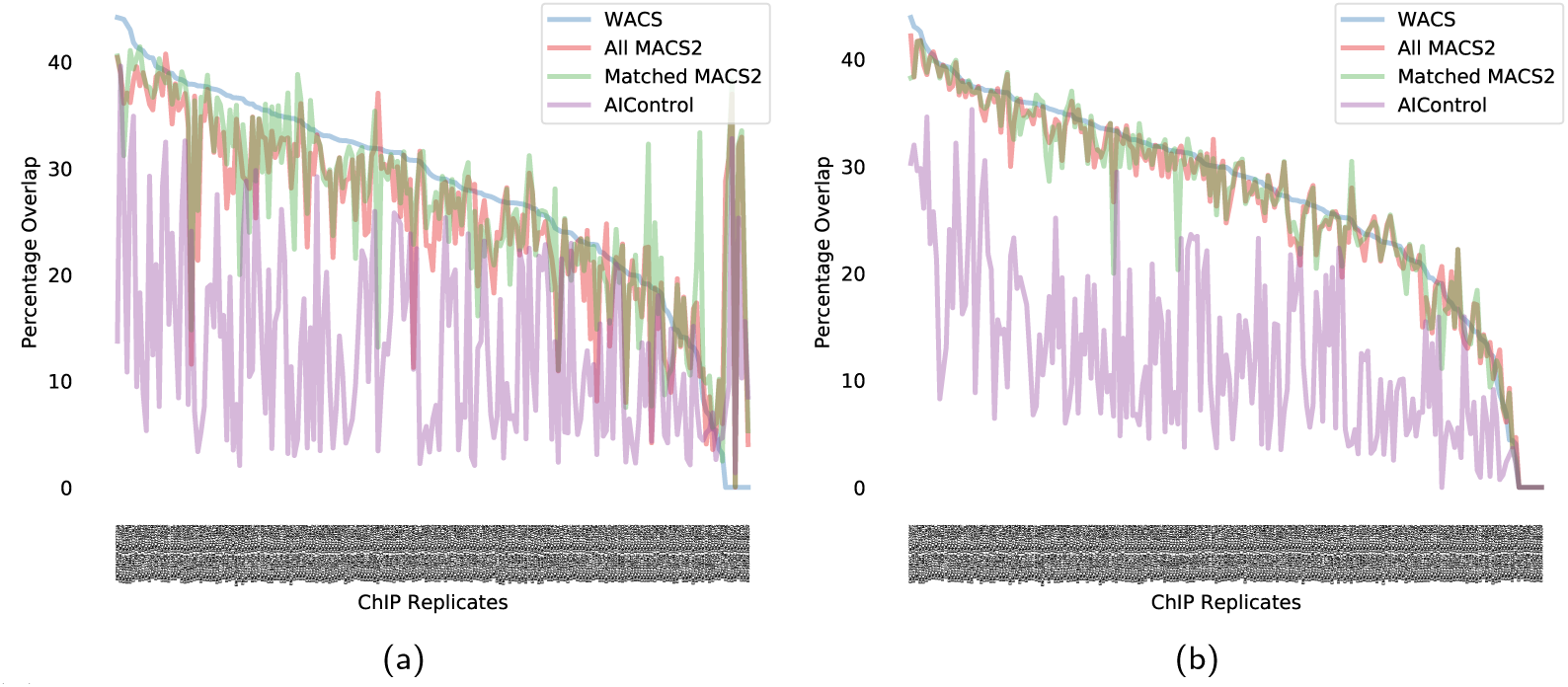
(a) Percentage overlap in all peaks between ENCODE replicates, for each of the four peak calling methods. (b) Percentage overlap in standardized peaks between ENCODE replicates, for each of the four peak calling methods

AIControl has the highest variability, in comparison to MACS2 and WACS, with the percentage overlap fluctuating between 5% and 34%. This shows poor reproducibility between replicates for AIControl. All MACS2 and Matched MACS2 have similar performance in terms of percentage overlap. The difference in percentage overlap between WACS and AIControl is noticeably much larger than the difference between WACS and MACS2 across the ChIP-seq replicates.

### Controls used per treatment sample

Our results (and other results [21, 22, 23]) for motif enrichment and reproducibility analysis suggest that smart controls offer superior background subtraction and peak-calling for ChIP-seq data. However, the standard practice remains to generate controls alongside each ChIP-seq experiment, or to match them on the basis of experimental details, such as cell/tissue type, read length and sequencer. If smart controls are to be used, it is unclear how many controls should be considered, and how many will end up in the smart control. It is unclear whether ENCODE matched controls are, in fact, the best choices or even among the controls selected by a smart control procedure.

Here, we aim to increase our understanding of the smart controls used to model the background signal. Figure 4 displays a matrix where the rows and columns represent the ChIP-seq and control datasets respectively. The blue color in the matrix represents the controls selected by WACS to fit each ChIP-seq dataset, the maroon color represents the ENCODE matched controls [24] and the magenta color represents the controls selected by both ENCODE and WACS.

**Figure 4:**
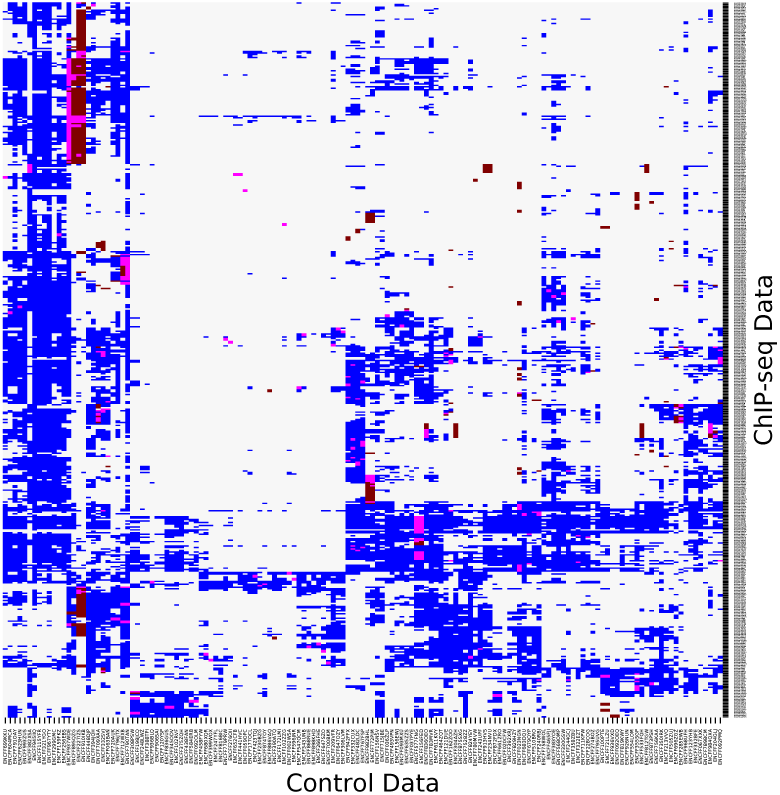
Comparison of controls used by WACS and ENCODE. The rows and columns correspond to the ChIP-seq and control experiments respectively. For each ChIP-seq dataset, the controls are given a *blue* color if they are used by WACS only, a *maroon* color if they are ENCODE matched controls only, and a *magenta* color if they are used by both ENCODE and WACS.

Let us first consider the WACS selected controls per ChIP-seq dataset (blue in Figure 4). Different subsets of the 147 controls are required by WACS for each ChIP-seq dataset, but these form several coherent clusters, where groups of ChIP-seq datasets use relatively the same controls for modeling the background signal. For example, the 10 or so controls most towards the left of the diagram are used in modeling nearly all the ChIP-seq, excepts those towards the bottom. The next 10 controls are widely used, though less so, and are distinct in be used for some of the ChIP-seqs towards the bottom. Conversely, there is a set of ChIP-seq datasets about 1/3 of the way from the bottom they rely on a large number of controls for modeling their background, whereas ChIP-seqs in the upper half rely almost solely on the leftmost controls. Although each ChIP-seq’s background is modeled by a unique combination of controls, a clear trend is that many controls are combined—approximately 31 on average, and up to 100 for some samples. Supplementary Figure 4 shows a histogram of the overall number of controls used by the ChIP-seq datasets using WACS.

For the ENCODE matched controls, we observe a range of 1 to 4 ENCODE matched controls per ChIP-seq dataset (maroon color in Figure 4). For 164 of the 438 ChIP-seq datasets (37%), none of the matched ENCODE controls are used to model the background signal in comparison to those used by WACS (rows with no magenta color in Figure 4). For example, 15 controls are used to model the background signal for the ChIP-seq dataset ENCFF025QKV in Figure 4, none of which are the matched ENCODE controls. For the remaining 63% of the ChIP-seq datasets, some of the ENCODE matched controls are also those selected by WACS, as seen in Figure 2 (magenta color). There are 207 ChIP-seq datasets (out of 438) that use all their matched ENCODE controls (in addition to other controls samples), and 32 ChIP-seq datasets that use half of their matched control samples to model the background signal.

### Validation on additional cell lines

Here, we further evaluate WACS, MACS2 and AIControl on three other cell lines: A549, HepG2 and GM12878. We specifically explored 20 ChIP-seq and 18 control datasets for each cell line. (See Supplementary Tables 4, 5 and 6 for accession codes of the samples.) We evaluated MACS2 with the ENCODE matched controls (Matched MACS2), MACS2 with the cell line specific controls (All MACS2), WACS with the cell line specific controls (WACS), WACS with the all controls across the three different cell lines (WACS AllCtrls), and AIControl with its predefined set of controls on ChIP-seq datasets (AIControl).

To evaluate the quality of the peaks generated by each method for each cell line, we first investigate motif enrichment. Figure 5 displays the motif enrichment for all and standardized peaks for each of the ChIP-experiments corresponding to each cell line, when using WACS (blue line), WACS AllCtrls (yellow line), All MACS2 (red line), Matched MACS2 (green line) and AIControl (purple line). AIControl across all cell lines, for all and standardized peaks, has the lowest motif enrichment. For the cell line A549, as seen in Figures 5a and 5d, WACS and WACS All Ctrls display the highest motif enrichment and have very similar performance. WACS and WACS All Ctrls outperform Matched MACS2, All MACS2 and AIControl on 14 treatment samples in total, as shown in Table 3. The differences for A549 with standardized peaks are statistically significant using a paired sign test and at a p-value of less than 10^−5^. An equivalent trend is observed for the GM12878 cell line (Figures 5b and 5e). However, when using all peaks, WACS has the highest motif enrichment; WACS outperforms WACS All Ctrls, Matched MACS2, All MACS2 and AIControl on 15 treatment samples in total, as shown in Table 3. The differences for GM12878, with both standard and all peaks, are statistically significant using a paired sign test and at a p-value of less than 10^−5^. Additionally, for standardized peaks, for cell lines A549 and GM12878, we notice almost equivalent motif enrichment when using All MACS2 and Matched MACS2. For HepG2 with all peaks (Figure 5c), on the other hand, Matched MACS2 outperforms WACS, WACS All Ctrls, All MACS2 and AIControl on 11 treatment samples in total. For HepG2 with standardized peaks (Figure 5f), all methods display similar performance. For HepG2, the differences between All MACS2 and WACS are statistically significant using a paired sign test and at a p-value of less than 10^−5^.

**Table 3:** Motif Enrichment Summary.

| Cell Line | WACS | WACS<br>AllCtrls | Matched<br>MACS2 | All<br>MACS2 | AIControl |
| --- | --- | --- | --- | --- | --- |
| **All Peaks** |  |  |  |  |  |
| A549 | 7 | 7 | 2 | 4 | 0 |
| GM12878 | 15 | 2 | 2 | 1 | 0 |
| HepG2 | 0 | 9 | 11 | 0 | 0 |
| **Standardized** |  |  |  |  |  |
| A549 | 10 | 8 | 1 | 1 | 0 |
| GM12878 | 11 | 5 | 3 | 0 | 1 |
| HepG2 | 3 | 6 | 5 | 0 | 6 |

**Figure 5:**
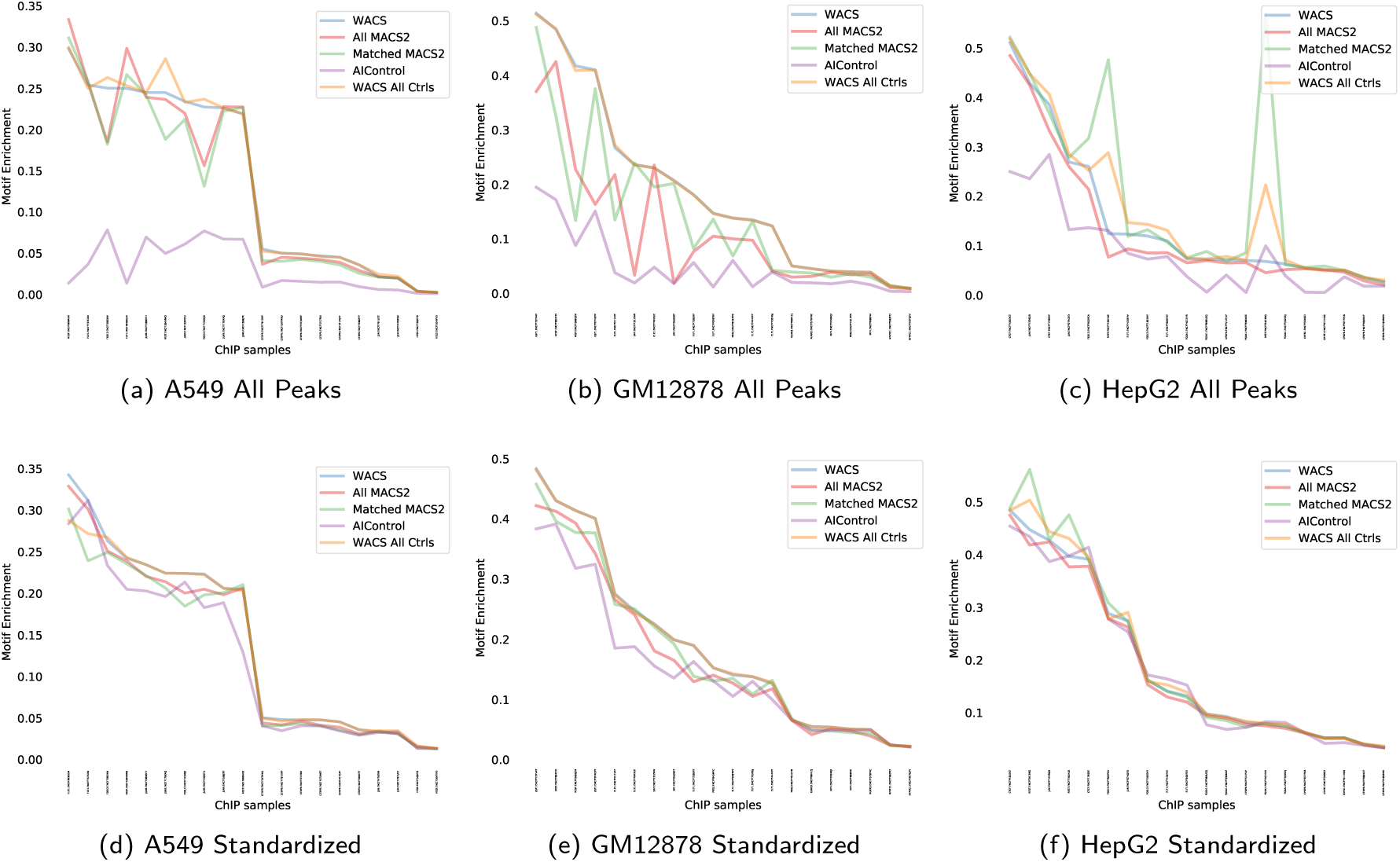
Motif enrichment of the treatment samples for each of the five peak calling methods for each of the 3 cell lines: A549 (*a* and *d*), GM12878 (*b* and *e*) and HepG2 (*c* and *f*).

Next, we explore the reproducibility of peaks in ChIP-seq replicates for each cell line. There are a total of 10 ChIP-seq experiments for each cell line, each with two replicates. Figure 6 show the percentage overlap with all and standardized peaks for each of the ChIP-seq experiments, when using WACS (blue line), WACS AllCtrls (yellow line), All MACS2 (red line), Matched MACS2 (green line) and AIControl (purple line). WACS All Ctrls outperforms WACS, Matched MACS2, All MACS2 and AIControl on all of the ChIP-seq datasets for all the three cell lines, A549, GM12878 and HepG2 for all and standardized peaks, as show in Table 4. Again, AIControl displays the lowest percentage overlap for A549, GM12878 and HepG2 for all and standardized peaks.

**Table 4:** Reproducibility Analysis Summary.

| Cell Line | WACS | WACS<br>AllCtrls | Matched<br>MACS2 | All<br>MACS2 | AIControl |
| --- | --- | --- | --- | --- | --- |
| **All Peaks** |  |  |  |  |  |
| A549 | 1 | 9 | 2 | 0 | 1 |
| GM12878 | 3 | 4 | 1 | 1 | 1 |
| HepG2 | 1 | 7 | 2 | 0 | 0 |
| **Standardized** |  |  |  |  |  |
| A549 | 2 | 9 | 2 | 0 | 0 |
| GM12878 | 2 | 4 | 3 | 1 | 0 |
| HepG2 | 1 | 8 | 1 | 0 | 0 |

**Figure 6:**
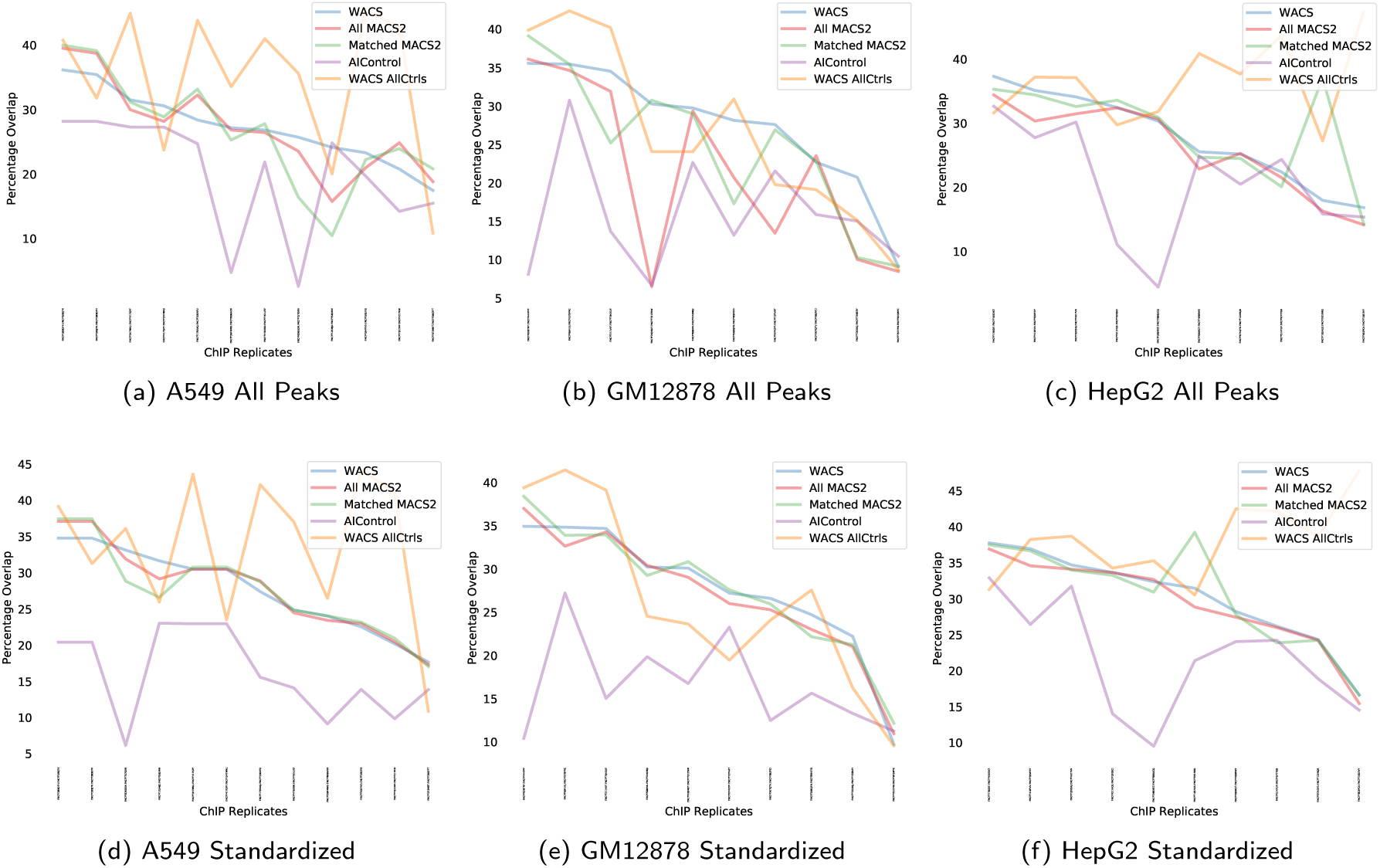
Percentage overlap in peaks between ENCODE replicates, for each of the five peak calling methods for each of the 3 cell lines: A549 (*a* and *d*), GM12878 (*b* and *e*) and HepG2 (*c* and *f*).

## Discussion

In this paper, we provide a method for improved peak-calling and increase our understanding of ChIP-seq data, controls and their biases. Firstly, we showed that the controls selected by WACS are not necessarily the matched ENCODE controls. Additionally, for most of the ChIP-seq datasets, many more than two controls are selected to model the background signal. This suggests that the ENCODE guidelines may need to be modified to include more than two “matched” controls per ChIP-seq dataset. We also notice that for a small number of controls, Matched MACS2 and WACS perform similarly. The more controls used, the better WACS performs. We showed that a form of intelligent control selection is beneficial for the combination (selection) of controls, as it better models the estimated background signal, where different controls represent different types of biases. As noted by Hiranuma et al. [23], this will also allow researchers to use other controls non-specific to their ChIP-seq experiment to model the noise distribution. This will decrease cost, time and resources required to perform the ChIP-seq experiments. WACS not only provides weights per control for a more efficient background model, it uses an already existing and precise peak calling method in MACS2.

Hiranuma et al. [23] claim that AIControl is better at removing background noise than MACS2. However, our results suggest the contrary (see Figure 5). This may be due to a number of reasons. First, Hiranuma et al. [23] uses a different and nonstandard evaluation method for reproducibility analysis. Whereas we adopted the widely used approach of looking at peak overlaps between biological replicates [4, 5], Hiranuma showed that AIControl had higher irreproducibility than MACS when applied to unrelated datasets. Furthermore, Hiranuma et al. applied MACS2 using only one matched control, while for our analysis, we used either all the ENCODE matched controls for a treatment sample or simply all controls from the same K562 cell line. In either case, the provision of multiple controls may have improved MACS2’s performance.

In this manuscript, we described using NNLS to fit a model of ChIP-seq background to control densities, but other formulations are possible. For example, we experimented with an instance-weighted NNLS formulation, to account for differing variances on the regression targets *y*_*i*_ (the ChIP-seq read counts per window). We did not find any improvement in performance. However, results may depend on how one estimates target variances. Relatedly, performing regression on log-transformed read counts may be worth exploring. RNA-seq analysis tools such as DESeq2 [33] use log linear models for read counts and comparisons between conditions. It would also make sense to explore L1-penalized regression formulations, to explore trade offs between the number of controls used to model background and the accuracy of the background model.

Future work will deal with a more thorough analysis of the weighted controls approach on other high throughput sequencing data, such as RNA-seq, and other cell lines. The weighted approach will be used to study the biases in RNA-seq data across different platforms, labs, cell types, tissues, etc. For example, RNA-seq is used to measure the difference in gene expression between tissues, where a tissue consists of a mixture of cell types. To generate a realistic control tissue, the weighted approach can be used to weight the cell types in the tissue to model the background signal. Also, in this analysis, we focused on sharp peaks, which are more generally found at protein-DNA binding sites. Thus, an analysis of other broader peaks, for example, will be conducted. Ultimately, the overall aim is to further our understanding of the significance of the increasing amount of publicly available data in peak calling analysis to obtain more efficient results.

## Conclusion

We developed a peak calling method, WACS, which allows a mixture of weighted controls as input. The user inputs the controls. These controls can either be weighted by the user, or the weights can be computed by our regression approach. The latter systematically estimates the weights of the input controls to model the background signal for that ChIP-seq experiment. In the special case of equal weights which sum up to 1, the peaks output from WACS and MACS2 are identical. If different weights are allowed, the two algorithms have different outputs. WACS allows only positive weights for better interpretability of results. Negative weights are biologically difficult to interpret; as it does not add to the background signal. WACS proceeds to use this devised background signal to identify regions of enrichment along the genome. WACS is an extension of the most highly cited peak calling algorithm, MACS2 [18]. We conducted a comparison between WACS, MACS2 and AIControl to evaluate our method and the significance of the weighted controls. WACS significantly outperforms both MACS2 and AIControl in motif enrichment analysis and reproducibility analysis.

## Ethics approval and consent to participate

Not applicable.

## Consent for publication

Not applicable.

## Competing interests

The authors declare that they have no competing interests.

## Availability of data and materials

ChIP-seq data used to develop and evaluate this method can be found online on the ENCODE website https://www.encodeproject.org. The WACS software can be found on the following website: https://www.perkinslab.ca/software.

## Funding

This work was supported in part by a Queen Elizabeth II Graduate Scholarship in Science and Technology (QEII-GSST) to AA, and by an NSERC Discovery Grant to TJP.

## Author’s contributions

AA and TJP conceived and designed the analysis. AA developed the tool, performed analysis/computations and wrote the manuscript with input from TJP. TJP edited the manuscript. TJP and MT supervised the project. All authors provided critical feedback and helped shape the research, analysis and manuscript.

## Acknowledgments

We thank Compute Canada for granting us access to their cluster to store data and run our computational analyses. We also thank members of the Perkins lab for their feedback.

## Additional Files

Additional file 1 — WACSSupp

Includes Supplementary Tables 1, 2, 3, 4, 5 and 6. Supplementary Figures 1, 2, 3 and 4.

