## Supplementary Information for "WACS: Improving ChIP-seq Peak Calling by Optimally Weighting Controls"

### Weighted analysis of ChIP-seq

Table 1: Table for the ChIP-seq experiments and their corresponding ChIP-seq replicate samples and TFs for the K562 cell line from the ENCODE database used in our analysis.

| Experiment | Replicates | TF |
|---|---|---|
| ENCSR440VKE | ENCFF540ZTB, ENCFF666EIZ | ADNP |
| ENCSR641BSL | ENCFF471TDC, ENCFF360QAB | AGO1 |
| ENCSR822CCM | ENCFF225IQS, ENCFF431WVX | ARID1B |
| ENCSR000EFY | ENCFF477DKP, ENCFF495MEU | ARID3A |
| ENCSR613NUC | ENCFF258DFY, ENCFF490XTX | ARNT |
| ENCSR155KHM | ENCFF928JGK, ENCFF108SDD | ARNT |
| ENCSR091GVJ | ENCFF774RUP, ENCFF892LAS | ATF1 |
| ENCSR869IUD | ENCFF719FYJ, ENCFF883MZR | ATF2 |
| ENCSR000BNU | ENCFF438DHS, ENCFF308RGN | ATF3 |
| ENCSR632DCH | ENCFF938FQC, ENCFF327XSI | ATF3 |
| ENCSR740NPG | ENCFF221XVM, ENCFF042SJF | BACH1 |
| ENCSR000EGD | ENCFF151GOI, ENCFF540DXB | BACH1 |
| ENCSR492LTS | ENCFF323IVG, ENCFF862XFS | BCLAF1 |
| ENCSR000BKH | ENCFF554KEG, ENCFF339YBR | BCLAF1 |
| ENCSR808AKZ | ENCFF207LND, ENCFF541XEW | BCOR |
| ENCSR000EGV | ENCFF213JOK, ENCFF988VSQ | BHLHE40 |
| ENCSR782WRO | ENCFF918CSI, ENCFF110FDC | BMI1 |
| ENCSR583ACG | ENCFF652HPU, ENCFF860JBC | BRD4 |
| ENCSR350XWY | ENCFF904OXK, ENCFF077NYA | C11orf30 |
| ENCSR948QLZ | ENCFF382DBE, ENCFF888AOR | CBX1 |
| ENCSR000BRT | ENCFF093QMN, ENCFF289RGM | CBX3 |
| ENCSR272JAT | ENCFF468LGZ, ENCFF092TPD | CBX5 |
| ENCSR000ATW | ENCFF781JEI, ENCFF954PRL | CBX8 |
| ENCSR121PFY | ENCFF201MXQ, ENCFF049RJI | CDC5L |
| ENCSR416QLJ | ENCFF002PNZ, ENCFF346SNT | CEBPB |

| | | |
|---|---|---|
| ENCSR000BRQ | ENCFF154YTN, ENCFF314OQP | CEBPB |
| ENCSR000EHE | ENCFF953ZZL, ENCFF729WGC | CEBPB |
| ENCSR620VIC | ENCFF693JVP, ENCFF620VNP | CEBPG |
| ENCSR618GDK | ENCFF799RPS, ENCFF576EDK | CEBPZ |
| ENCSR065XVO | ENCFF432NAV, ENCFF824HGA | CHAMP1 |
| ENCSR130HEG | ENCFF423MBY, ENCFF307UCL | COPS2 |
| ENCSR109YGM | ENCFF358SSM, ENCFF405YCF | CREB3L1 |
| ENCSR093FKD | ENCFF294DBF, ENCFF334OKF | CREB3 |
| ENCSR077DKV | ENCFF363YKW, ENCFF473NMO | CREM |
| ENCSR201NQZ | ENCFF331BYU, ENCFF809OVE | CTBP1 |
| ENCSR000BNK | ENCFF454MMY, ENCFF310MOR | CTCFL |
| ENCSR000BPJ | ENCFF691BQZ, ENCFF391HFU | CTCF |
| ENCSR000DWE | ENCFF081HVQ, ENCFF494VZW | CTCF |
| ENCSR000AKO | ENCFF871GXE, ENCFF550BMQ | CTCF |
| ENCSR000EGM | ENCFF198CVB, ENCFF488CXC | CTCF |
| ENCSR178NTX | ENCFF589UVV, ENCFF060WJH | CUX1 |
| ENCSR446LAV | ENCFF546ZLD, ENCFF562ATM | DDX20 |
| ENCSR382AIB | ENCFF810IRF, ENCFF760JNE | DDX20 |
| ENCSR387SYS | ENCFF056XVW, ENCFF540SXT | DEAF1 |
| ENCSR167JBG | ENCFF384DMV, ENCFF730MAX | DIDO1 |
| ENCSR987PBI | ENCFF573OTF, ENCFF378UVV | DNMT1 |
| ENCSR715CCR | ENCFF990RGE, ENCFF205JWI | DPF2 |
| ENCSR030OKC | ENCFF569XLF, ENCFF824KAR | DROSHA |
| ENCSR563LLO | ENCFF846CYU, ENCFF226KOW | E2F1 |
| ENCSR000EWL | ENCFF613CDR, ENCFF156NIH | E2F4 |
| ENCSR709DRM | ENCFF980YNA, ENCFF564FBJ | E2F5 |
| ENCSR000EWJ | ENCFF201BQU, ENCFF011OSI | E2F6 |
| ENCSR000BLI | ENCFF823GCX, ENCFF827SLL | E2F6 |
| ENCSR000BMD | ENCFF331TRC, ENCFF535PLW | ELF1 |
| ENCSR975SSR | ENCFF251EVS, ENCFF619SRD | ELF1 |
| ENCSR502OEK | ENCFF477CYZ, ENCFF784MTZ | ELF1 |
| ENCSR638QHV | ENCFF958WMW, ENCFF339RWU | ELF4 |
| ENCSR338QAC | ENCFF407GAB, ENCFF137QTZ | ELK1 |
| ENCSR000EFU | ENCFF865ISL, ENCFF554BLC | ELK1 |
| ENCSR000EGE | ENCFF982AFE, ENCFF200PYZ | EP300 |
| ENCSR486IFJ | ENCFF857YYV, ENCFF320WXN | ESRRA |
| ENCSR000BKQ | ENCFF401KIO, ENCFF398SXO | ETS1 |

| | | |
|---|---|---|
| ENCSR596IKD | ENCFF950PAH, ENCFF535SSL | ETS2 |
| ENCSR277DMR | ENCFF656BKZ, ENCFF670SFK | ETV1 |
| ENCSR000FCE | ENCFF002FHN, ENCFF836NST | ETV6 |
| ENCSR124BJR | ENCFF417FHN, ENCFF003FNG | ETV6 |
| ENCSR159OCC | ENCFF515YFJ, ENCFF780DBH | FLAG-ATF1 |
| ENCSR633EIC | ENCFF072SKI, ENCFF030XLK | FLAG-PBX2 |
| ENCSR239ZLZ | ENCFF581AZU, ENCFF500HFO | FOSL1 |
| ENCSR000BMV | ENCFF492NUF, ENCFF561ILM | FOSL1 |
| ENCSR819LHG | ENCFF364SJK, ENCFF611PXX | FOXA1 |
| ENCSR847LBF | ENCFF834YLT, ENCFF904PBR | FOXJ2 |
| ENCSR302AWT | ENCFF703CKR, ENCFF931HHQ | FOXK2 |
| ENCSR508DQA | ENCFF563FZR, ENCFF373AFZ | FOXK2 |
| ENCSR429QPP | ENCFF182YDN, ENCFF855NIZ | FOXM1 |
| ENCSR051DXE | ENCFF361MTW, ENCFF503JQK | FUS |
| ENCSR290MUH | ENCFF400MMA, ENCFF593XTM | GABPA |
| ENCSR000EWM | ENCFF642RHD, ENCFF788ICY | GATA1 |
| ENCSR000EWG | ENCFF340JWK, ENCFF779CRV | GATA2 |
| ENCSR000DKA | ENCFF880ZXO, ENCFF082TOF | GATA2 |
| ENCSR801RPW | ENCFF709ZMI, ENCFF438XEX | GTF2A2 |
| ENCSR532KTI | ENCFF623YPH, ENCFF106BHY | GTF2E2 |
| ENCSR000EHC | ENCFF215TDO, ENCFF040MZK | GTF2F1 |
| ENCSR000APC | ENCFF472XNE, ENCFF407AUC | H2AFZ |
| ENCSR000AKP | ENCFF301TVL, ENCFF879BWC | H3K27ac |
| ENCSR000AKQ | ENCFF190OWE, ENCFF692KQZ | H3K27me3 |
| ENCSR000EWB | ENCFF915XIL, ENCFF330YFF | H3K27me3 |
| ENCSR000AKR | ENCFF639PLN, ENCFF673KBG | H3K36me3 |
| ENCSR000DWB | ENCFF975JFV, ENCFF989ORU | H3K36me3 |
| ENCSR000EWC | ENCFF290LQY, ENCFF063EDR | H3K4me1 |
| ENCSR000AKT | ENCFF010SKB, ENCFF773VGC | H3K4me2 |
| ENCSR000DWD | ENCFF185YRK, ENCFF955AMI | H3K4me3 |
| ENCSR000EWA | ENCFF706SCF, ENCFF611YPB, ENCFF792ZLC, ENCFF742FDS | H3K4me3 |
| ENCSR668LDD | ENCFF236SNL, ENCFF661UGK | H3K4me3 |
| ENCSR000AKU | ENCFF633WWH, ENCFF777LZD | H3K4me3 |
| ENCSR000APD | ENCFF408YHI, ENCFF947DVY | H3K79me2 |
| ENCSR000AKV | ENCFF103YPC, ENCFF698ROL | H3K9ac |
| ENCSR000EVZ | ENCFF159SIY, ENCFF869PBP, ENCFF763ZGN, ENCFF236CJR | H3K9ac |

| | | |
|---|---|---|
| ENCSR000APE | ENCFF146NLP, ENCFF559DHZ | H3K9me3 |
| ENCSR000EFN | ENCFF068XVC, ENCFF976DWY | HCFC1 |
| ENCSR568PGX | ENCFF785NOU, ENCFF135YTU | HDAC1 |
| ENCSR387UWP | ENCFF677JTW, ENCFF921IAQ | HDAC1 |
| ENCSR000AQG | ENCFF456VOS, ENCFF610YXS | HDAC2 |
| ENCSR075HTM | ENCFF408BKE, ENCFF497GOO | HDAC2 |
| ENCSR000ATJ | ENCFF213CZL, ENCFF428XVO | HDAC6 |
| ENCSR000DJZ | ENCFF786TLS, ENCFF659KOL | HDAC8 |
| ENCSR835TCD | ENCFF997BDJ, ENCFF285FGV | HDAC8 |
| ENCSR563YDA | ENCFF769DDY, ENCFF221CPW | HDGF |
| ENCSR197ALX | ENCFF702WHX, ENCFF795OAZ | HDGF |
| ENCSR091JXL | ENCFF502NAY, ENCFF046ACV | HES1 |
| ENCSR619GFP | ENCFF448SJG, ENCFF786KLU | HINFP |
| ENCSR757IIU | ENCFF797FVU, ENCFF202BLM | HMBOX1 |
| ENCSR296MXW | ENCFF733RTM, ENCFF275XJW | HNRNPUL1 |
| ENCSR005NMT | ENCFF695GTB, ENCFF115BME | ID3 |
| ENCSR948VFL | ENCFF016KRJ, ENCFF880SVB | IKZF1 |
| ENCSR395HWC | ENCFF013MXK, ENCFF752FIR | IKZF1 |
| ENCSR648INT | ENCFF145ZDW, ENCFF831ODM | ILK |
| ENCSR000EGT | ENCFF489YJG, ENCFF728WOA | IRF1 |
| ENCSR000EGK | ENCFF168KTM, ENCFF299HHL | IRF1 |
| ENCSR000EGU | ENCFF218JFL, ENCFF158ZVB | IRF1 |
| ENCSR854MCV | ENCFF922JWO, ENCFF701KAF | IRF1 |
| ENCSR000EGL | ENCFF259RRK, ENCFF244IJF | IRF1 |
| ENCSR376WCJ | ENCFF178EJY, ENCFF057KBN | IRF2 |
| ENCSR926KTP | ENCFF623MQU, ENCFF634KQT | IRF9 |
| ENCSR795IYP | ENCFF512YJL, ENCFF090SFR | JUNB |
| ENCSR000EGN | ENCFF400BSN, ENCFF321ZQU | JUND |
| ENCSR000EZX | ENCFF479JUJ, ENCFF643UCP | JUN |
| ENCSR000EFS | ENCFF924CYX, ENCFF014UUB | JUN |
| ENCSR000EZW | ENCFF814CHV, ENCFF050LIC | JUN |
| ENCSR000EZT | ENCFF749RRI, ENCFF784MLU | JUN |
| ENCSR000EGH | ENCFF703YNU, ENCFF527CTN | JUN |
| ENCSR086FZL | ENCFF578CKQ, ENCFF707JIA | KAT8 |
| ENCSR360HRA | ENCFF042LKP, ENCFF342FIU | KDM1A |
| ENCSR908CMW | ENCFF008IZS, ENCFF599IYT | KDM1A |
| ENCSR642VZY | ENCFF330ZVF, ENCFF066TDG | KDM4B |

| | | |
|---|---|---|
| ENCSR608HVP | ENCFF829XLC, ENCFF771KEQ | KLF13 |
| ENCSR760UVO | ENCFF749LRJ, ENCFF408TSG | KLF16 |
| ENCSR550HCT | ENCFF378QWS, ENCFF878UKW | KLF1 |
| ENCSR735JCD | ENCFF966VXD, ENCFF370NVV | KMT2B |
| ENCSR530XQI | ENCFF799FNR, ENCFF856TBD | L3MBTL2 |
| ENCSR343ELW | ENCFF933HFV, ENCFF178YFO | LEF1 |
| ENCSR000EGI | ENCFF589IXE, ENCFF097GYE | MAFF |
| ENCSR818DQV | ENCFF253HOC, ENCFF773DBT | MAFG |
| ENCSR000EGX | ENCFF719OIE, ENCFF852HSQ | MAFK |
| ENCSR000EFV | ENCFF635HIS, ENCFF360QBV | MAX |
| ENCSR221GAN | ENCFF502VUK, ENCFF655OEX | MBD2 |
| ENCSR990AZC | ENCFF668FCG, ENCFF145NON | MCM3 |
| ENCSR079WHK | ENCFF125HKL, ENCFF046VNU | MCM5 |
| ENCSR038RGL | ENCFF245YPG, ENCFF980TTL | MCM7 |
| ENCSR542WJU | ENCFF548AHK, ENCFF995UTD | MCM7 |
| ENCSR000BNV | ENCFF850FRG, ENCFF401BQM | MEF2A |
| ENCSR647ZXA | ENCFF047IWR, ENCFF662HRL | MEF2D |
| ENCSR710WLO | ENCFF066JMV, ENCFF330XAC | MGA |
| ENCSR426MDV | ENCFF639CLN, ENCFF682OCC | MIER1 |
| ENCSR000FCB | ENCFF677BBS, ENCFF983RUX | MITF |
| ENCSR675LRO | ENCFF353XMB, ENCFF131XTL | MLLT1 |
| ENCSR107GRP | ENCFF887LAD, ENCFF777ZPZ | MLLT1 |
| ENCSR390VGH | ENCFF803TYM, ENCFF698VVG | MNT |
| ENCSR979QYJ | ENCFF036IOO, ENCFF358HOW | MNT |
| ENCSR113LAS | ENCFF254IUH, ENCFF167NVK | MTA2 |
| ENCSR411UYA | ENCFF563DGZ, ENCFF750EKT | MTA2 |
| ENCSR000EGZ | ENCFF726UVN, ENCFF388QNA | MXI1 |
| ENCSR000EGJ | ENCFF641HZR, ENCFF006GQZ | MYC |
| ENCSR000EGS | ENCFF239WGU, ENCFF836IMK | MYC |
| ENCSR000FAZ | ENCFF439NNR, ENCFF483RLD | MYC |
| ENCSR000EZV | ENCFF357MHM, ENCFF491EUN | MYC |
| ENCSR000EZU | ENCFF014DRH, ENCFF553WMU | MYC |
| ENCSR737LTZ | ENCFF179DZP, ENCFF022IWG | MYNN |
| ENCSR085QEV | ENCFF397LNB, ENCFF700WSI | NBN |
| ENCSR931HNY | ENCFF427CAA, ENCFF314PZG | NCOA1 |
| ENCSR910JAI | ENCFF420DGT, ENCFF129RYE | NCOR1 |
| ENCSR298JCG | ENCFF247FGI, ENCFF488BQX | NCOR1 |

| | | |
|---|---|---|
| ENCSR798ILC | ENCFF072KEJ, ENCFF033JFM | NCOR1 |
| ENCSR632SHZ | ENCFF027QPB, ENCFF697TDR | NFE2L1 |
| ENCSR000FCC | ENCFF198RTF, ENCFF187ZET | NFE2 |
| ENCSR552YGL | ENCFF945TUI, ENCFF581SOT | NFE2 |
| ENCSR657EOF | ENCFF530UEA, ENCFF437UWB | NFRKB |
| ENCSR085DDI | ENCFF833ZOK, ENCFF318SPV | NFXL1 |
| ENCSR415TXN | ENCFF203MUO, ENCFF618MHM | NONO |
| ENCSR886RYH | ENCFF008CUX, ENCFF549MBS | NONO |
| ENCSR178DEG | ENCFF285CII, ENCFF096KBP | NR2C1 |
| ENCSR742IDN | ENCFF149WGG, ENCFF315DCZ | NR2C1 |
| ENCSR000EWH | ENCFF771NSF, ENCFF078BWN | NR2C2 |
| ENCSR750LYM | ENCFF597WPX, ENCFF856EXS | NR2C2 |
| ENCSR970NKQ | ENCFF550IUC, ENCFF284XYU | NR2F1 |
| ENCSR000BRS | ENCFF568RDF, ENCFF665EXC | NR2F2 |
| ENCSR707QWA | ENCFF908MXL, ENCFF507UJN | NR2F6 |
| ENCSR000DJW | ENCFF602UAC, ENCFF650RFL | NR4A1 |
| ENCSR130PDE | ENCFF908PSF, ENCFF786GEU | NR4A1 |
| ENCSR837EYC | ENCFF564KCD, ENCFF207NLX | NRF1 |
| ENCSR494TDU | ENCFF337GMY, ENCFF821HMU | NRF1 |
| ENCSR998AJK | ENCFF564CXM, ENCFF722LJA | NRF1 |
| ENCSR263DFP | ENCFF616QUW, ENCFF577WGL | PBX2 |
| ENCSR924GXX | ENCFF229VMU, ENCFF802CIM | PHB2 |
| ENCSR000AQH | ENCFF509IQQ, ENCFF302WSZ | PHF8 |
| ENCSR115SMW | ENCFF494OJC, ENCFF636IGK | PKNOX1 |
| ENCSR000FAY | ENCFF976EMR, ENCFF323MTY | POLR2A |
| ENCSR031TFS | ENCFF559BJS, ENCFF725WJS | POLR2A |
| ENCSR000FAW | ENCFF191CWT, ENCFF536TPK | POLR2A |
| ENCSR388QZF | ENCFF438GBD, ENCFF601CVJ | POLR2A |
| ENCSR000BMR | ENCFF569DZM, ENCFF140UJB | POLR2A |
| ENCSR000EHP | ENCFF548NQU, ENCFF914ABF | POLR2A |
| ENCSR000EGF | ENCFF630JRN, ENCFF594OID | POLR2A |
| ENCSR400FSM | ENCFF241ZAF, ENCFF825VBC | POLR2H |
| ENCSR220YXI | ENCFF367OYT, ENCFF895DLK | PRPF4 |
| ENCSR948KMB | ENCFF525JGC, ENCFF031YZL | PTBP1 |
| ENCSR126FZN | ENCFF400SKV, ENCFF498KCA | PTRF |
| ENCSR314BBS | ENCFF586KRI, ENCFF662KUU | PTTG1 |
| ENCSR410DWC | ENCFF434YXL, ENCFF649ORS | PYGO2 |

| | | |
|---|---|---|
| ENCSR524BUE | ENCFF168TBA, ENCFF024NZQ | RAD51 |
| ENCSR670JDQ | ENCFF459MNI, ENCFF706XFF | RB1 |
| ENCSR822LBD | ENCFF161VHM, ENCFF156OFZ | RBFOX2 |
| ENCSR772EEN | ENCFF158BYF, ENCFF557QLG | RELA |
| ENCSR000BMW | ENCFF204TUQ, ENCFF546IMN | REST |
| ENCSR137ZMQ | ENCFF268CBK, ENCFF253JTJ | REST |
| ENCSR968GIB | ENCFF837ZOY, ENCFF183MSQ | RFX1 |
| ENCSR041AXL | ENCFF871KNP, ENCFF490ICL | RFX1 |
| ENCSR530IUH | ENCFF157HDL, ENCFF668VFP | RING1 |
| ENCSR820GND | ENCFF015UIX, ENCFF587HZM | RNF2 |
| ENCSR076YPO | ENCFF719XQY, ENCFF544DDY | RNF2 |
| ENCSR608XTF | ENCFF779IQL, ENCFF257OOQ | RNF2 |
| ENCSR138FUZ | ENCFF880ODJ, ENCFF726AYE | RNF2 |
| ENCSR588AKU | ENCFF011XRF, ENCFF812KIP | RUNX1 |
| ENCSR414TYY | ENCFF548STV, ENCFF350NUJ | RUNX1 |
| ENCSR000AQJ | ENCFF416LJL, ENCFF961HAZ | SAP30 |
| ENCSR000EWI | ENCFF728IYF, ENCFF874YYW | SETDB1 |
| ENCSR000BLR | ENCFF329LWM, ENCFF334QAB | SIN3A |
| ENCSR920BLG | ENCFF385PRZ, ENCFF315DZX | SIN3A |
| ENCSR657JLK | ENCFF323OVV, ENCFF050VHP | SIN3B |
| ENCSR000BNW | ENCFF331BZT, ENCFF584HKJ | SIX5 |
| ENCSR189PYJ | ENCFF257QQX, ENCFF925CYF | SMAD2 |
| ENCSR000FCD | ENCFF412DRO, ENCFF033UID | SMAD5 |
| ENCSR643VTW | ENCFF157HYJ, ENCFF606LUD | SMARCA4 |
| ENCSR587OQL | ENCFF350FCE, ENCFF744TCZ | SMARCA4 |
| ENCSR895HSJ | ENCFF122TGF, ENCFF914TEL | SMARCA5 |
| ENCSR157TCS | ENCFF294UTM, ENCFF387BDW | SMARCE1 |
| ENCSR000EGW | ENCFF355PQK, ENCFF365BCR | SMC3 |
| ENCSR754ZHU | ENCFF157WFG, ENCFF372NDR | SNRNP70 |
| ENCSR991ELG | ENCFF487SUP, ENCFF758LME | SP1 |
| ENCSR815ZDS | ENCFF623PVZ, ENCFF405EME | SREBF1 |
| ENCSR000FAV | ENCFF463LMJ, ENCFF403WNR | STAT1 |
| ENCSR000FAU | ENCFF089XMI, ENCFF263PLG | STAT1 |
| ENCSR000FAT | ENCFF615CWW, ENCFF940QAP | STAT2 |
| ENCSR000FBC | ENCFF951ITB, ENCFF876GQO | STAT2 |
| ENCSR894CGX | ENCFF079TBV, ENCFF772XIR | SUPT5H |
| ENCSR412CTM | ENCFF986KSB, ENCFF614RBF | SUZ12 |

| | | |
|---|---|---|
| ENCSR047LSJ | ENCFF641OBF, ENCFF408GAN | TAF15 |
| ENCSR000BKS | ENCFF025QKV, ENCFF919HAZ | TAF1 |
| ENCSR000BNM | ENCFF518PIS, ENCFF126SXE | TAF7 |
| ENCSR671GFC | ENCFF931HWG, ENCFF562PEW | TAF7 |
| ENCSR106FRG | ENCFF525CPA, ENCFF695TZQ | TAL1 |
| ENCSR429XTR | ENCFF682NGE, ENCFF466OLG | TARDBP |
| ENCSR033VAZ | ENCFF860IRS, ENCFF263MBV | TARDBP |
| ENCSR353HEP | ENCFF283ZIL, ENCFF903JXA | TARDBP |
| ENCSR000EGA | ENCFF964COC, ENCFF946HWT | TBL1XR1 |
| ENCSR000EGB | ENCFF188QGH, ENCFF719NDH | TBL1XR1 |
| ENCSR744WOO | ENCFF715AOC, ENCFF516TOF | TCF12 |
| ENCSR189TRZ | ENCFF283SLJ, ENCFF532BVM | TCF12 |
| ENCSR888XZK | ENCFF600TNR, ENCFF425MJK | TCF7L2 |
| ENCSR863KUB | ENCFF587PVQ, ENCFF742TAS | TCF7 |
| ENCSR635GTR | ENCFF949ENV, ENCFF280AEF | TEAD2 |
| ENCSR000BRK | ENCFF790FNF, ENCFF516UTV | TEAD4 |
| ENCSR017GBO | ENCFF070MZX, ENCFF795XBH | TFDP1 |
| ENCSR224IKA | ENCFF684CFB, ENCFF780GMX | TFDP1 |
| ENCSR000BNN | ENCFF393ZAK, ENCFF680COV | THAP1 |
| ENCSR871TKJ | ENCFF406TJQ, ENCFF446HGS | THRAP3 |
| ENCSR907MZR | ENCFF923LMN, ENCFF551IFV | TRIM24 |
| ENCSR957LDM | ENCFF079SPF, ENCFF738QLQ | TRIM24 |
| ENCSR000EVY | ENCFF166JNV, ENCFF921KTI, ENCFF170NGB, ENCFF426DLJ | TRIM28 |
| ENCSR000BRW | ENCFF904CPI, ENCFF397ZVS, ENCFF206YYH, ENCFF793KZB | TRIM28 |
| ENCSR787RVK | ENCFF498EFE, ENCFF302OZO | TSC22D4 |
| ENCSR690GUG | ENCFF010TVY, ENCFF057SMG | U2AF1 |
| ENCSR479QAJ | ENCFF762TAN, ENCFF401PIJ | U2AF2 |
| ENCSR000BKT | ENCFF794ABP, ENCFF587WWS | USF1 |
| ENCSR000EHG | ENCFF420ZVF, ENCFF434RFX | USF2 |
| ENCSR359NFW | ENCFF979BFW, ENCFF102ZOS | USF2 |
| ENCSR107RHZ | ENCFF715AMR, ENCFF195WOK | YBX1 |
| ENCSR567JEU | ENCFF954HWS, ENCFF124RUL | YBX3 |
| ENCSR000BMH | ENCFF567GMS, ENCFF067TWQ | YY1 |
| ENCSR000BKU | ENCFF044TAL, ENCFF651HPM | YY1 |
| ENCSR000EWF | ENCFF633CMO, ENCFF139WUV | YY1 |
| ENCSR286PCG | ENCFF261BXO, ENCFF035GBL | ZBED1 |

| | | |
|---|---|---|
| ENCSR331GDC | ENCFF290MEZ, ENCFF544ZPF | ZBTB11 |
| ENCSR706BJO | ENCFF625AJO, ENCFF212PQL | ZBTB11 |
| ENCSR230PTV | ENCFF555QMG, ENCFF273NVY | ZBTB2 |
| ENCSR876GXA | ENCFF632WSK, ENCFF226IAA | ZBTB33 |
| ENCSR237VLT | ENCFF873BRB, ENCFF149LTH | ZBTB40 |
| ENCSR000BME | ENCFF784SCN, ENCFF778PDI | ZBTB7A |
| ENCSR322CFO | ENCFF500YYZ, ENCFF717OYW | ZEB2 |
| ENCSR004GKA | ENCFF754GYS, ENCFF035CJB | ZEB2 |
| ENCSR920ASP | ENCFF680KUB, ENCFF757TIT | ZFX |
| ENCSR557RVF | ENCFF723QZO, ENCFF019PLG | ZHX1 |
| ENCSR882ERE | ENCFF071KOM, ENCFF236ZUN | ZKSCAN1 |
| ENCSR448UKK | ENCFF811CBJ, ENCFF413VAR | ZKSCAN8 |
| ENCSR907JPB | ENCFF384WCY, ENCFF417HNM | ZMIZ1 |
| ENCSR102KIN | ENCFF056HVT, ENCFF900QPQ | ZMYM3 |
| ENCSR000EGP | ENCFF682XGC, ENCFF975ZXB | ZNF143 |
| ENCSR011PEI | ENCFF540VVE, ENCFF503IYW | ZNF175 |
| ENCSR580IAO | ENCFF687QLR, ENCFF191EJM | ZNF197 |
| ENCSR099NCH | ENCFF205VDU, ENCFF109OWW | ZNF24 |
| ENCSR695EQB | ENCFF657TZQ, ENCFF738AVH | ZNF24 |
| ENCSR117WTM | ENCFF396AXY, ENCFF391BBO | ZNF24 |
| ENCSR385AHH | ENCFF686ZIH, ENCFF786MHJ | ZNF24 |
| ENCSR000EWN | ENCFF891GFG, ENCFF094FZQ | ZNF263 |
| ENCSR000EVX | ENCFF444ZUH, ENCFF474ETY | ZNF274 |
| ENCSR000EWE | ENCFF917UQJ, ENCFF419SAY | ZNF274 |
| ENCSR200JYP | ENCFF303ICH, ENCFF041ZVD | ZNF316 |
| ENCSR167KBO | ENCFF267EEY, ENCFF908CYX | ZNF316 |
| ENCSR334HSW | ENCFF373YTD, ENCFF304KSV | ZNF318 |
| ENCSR352BJL | ENCFF019RWB, ENCFF353EFN | ZNF318 |
| ENCSR674SCQ | ENCFF103HTZ, ENCFF864BUV | ZNF354B |
| ENCSR000EFP | ENCFF140ZMN, ENCFF171VBC | ZNF384 |
| ENCSR011NOZ | ENCFF362TFV, ENCFF007ALP | ZNF407 |
| ENCSR598TIR | ENCFF772UWJ, ENCFF606TJM | ZNF507 |
| ENCSR591CCL | ENCFF300AUR, ENCFF996AFH | ZNF512 |
| ENCSR149ZBI | ENCFF057CNY, ENCFF216TEN | ZNF584 |
| ENCSR603XLW | ENCFF284MLL, ENCFF465ZZG | ZNF589 |
| ENCSR845BCL | ENCFF433SUG, ENCFF461OAN | ZNF639 |
| ENCSR729HVR | ENCFF095OEN, ENCFF334BPF | ZNF644 |

| | | |
|---|---|---|
| ENCSR737UST | ENCFF910XWG, ENCFF338MSY | ZNF740 |
| ENCSR532EMP | ENCFF165STC, ENCFF694KHP | ZNF740 |
| ENCSR194IJN | ENCFF227TEG, ENCFF398UYF | ZNF766 |
| ENCSR257XVY | ENCFF690YWW, ENCFF180ZQP | ZNF83 |
| ENCSR175SZH | ENCFF122RSM, ENCFF450TNG | ZSCAN29 |
| ENCSR780BBJ | ENCFF548YYR, ENCFF566EXS | ZZZ3 |

Table 2: Table for the ChIP-seq samples and their corresponding control samples for the K562 cell line from the ENCODE database used in our analysis.

| ChIP-seq | Controls |
|---|---|
| ENCFF408TSG | ENCFF162ZOO, ENCFF092PMQ |
| ENCFF258DFY | ENCFF895QZG, ENCFF227IZS, ENCFF910IKB, ENCFF937WDE |
| ENCFF772UWJ | ENCFF230BZG |
| ENCFF110FDC | ENCFF156FED, ENCFF577FNG |
| ENCFF334BPF | ENCFF493VPN |
| ENCFF551IFV | ENCFF227IZS, ENCFF910IKB |
| ENCFF666EIZ | ENCFF609HLC |
| ENCFF002FHN | ENCFF332SVJ, ENCFF204GLJ |
| ENCFF122RSM | ENCFF227IZS, ENCFF910IKB |
| ENCFF330ZVF | ENCFF162ZOO, ENCFF332SVJ |
| ENCFF908CYX | ENCFF227IZS, ENCFF910IKB |
| ENCFF623MQU | ENCFF647SZO |
| ENCFF921IAQ | ENCFF895QZG, ENCFF227IZS, ENCFF910IKB, ENCFF937WDE |
| ENCFF719FYJ | ENCFF895QZG, ENCFF227IZS, ENCFF910IKB, ENCFF937WDE |
| ENCFF331BYU | ENCFF895QZG, ENCFF227IZS, ENCFF910IKB, ENCFF937WDE |
| ENCFF569XLF | ENCFF584ERB, ENCFF092OLM |
| ENCFF166JNV | ENCFF355SGP |
| ENCFF548AHK | ENCFF227IZS, ENCFF910IKB |
| ENCFF241ZAF | ENCFF448DJP |
| ENCFF283SLJ | ENCFF227IZS, ENCFF910IKB |
| ENCFF516TOF | ENCFF227IZS, ENCFF910IKB |
| ENCFF733RTM | ENCFF308PSW, ENCFF198JCQ |
| ENCFF996AFH | ENCFF752BZZ |
| ENCFF314PZG | ENCFF895QZG, ENCFF227IZS, ENCFF910IKB, ENCFF937WDE |
| ENCFF310MOR | ENCFF772PJM, ENCFF982BHL |
| ENCFF263PLG | ENCFF332CUX |

| | |
|---|---|
| ENCFF057CNY | ENCFF299GMC |
| ENCFF682XGC | ENCFF023NGN |
| ENCFF405YCF | ENCFF895QZG, ENCFF227IZS, ENCFF910IKB, ENCFF937WDE |
| ENCFF109OWW | ENCFF895QZG, ENCFF227IZS, ENCFF910IKB, ENCFF937WDE |
| ENCFF824HGA | ENCFF712WXB, ENCFF790TAN |
| ENCFF980TTL | ENCFF227IZS, ENCFF910IKB |
| ENCFF600TNR | ENCFF156FED, ENCFF577FNG |
| ENCFF398UYF | ENCFF321YGO |
| ENCFF887LAD | ENCFF895QZG, ENCFF227IZS, ENCFF910IKB, ENCFF937WDE |
| ENCFF363YKW | ENCFF709XAA, ENCFF332SVJ |
| ENCFF095OEN | ENCFF493VPN |
| ENCFF682OCC | ENCFF895QZG, ENCFF227IZS, ENCFF910IKB, ENCFF937WDE |
| ENCFF290MEZ | ENCFF213VVQ, ENCFF063BAN |
| ENCFF245YPG | ENCFF227IZS, ENCFF910IKB |
| ENCFF438DHS | ENCFF772PJM, ENCFF982BHL |
| ENCFF719NDH | ENCFF023NGN |
| ENCFF578CKQ | ENCFF227IZS, ENCFF910IKB |
| ENCFF042SJF | ENCFF274BOZ |
| ENCFF206YYH | ENCFF984QXA, ENCFF836OEO, ENCFF204GLJ, ENCFF304AZH, ENCFF533FQH |
| ENCFF950PAH | ENCFF816BIC |
| ENCFF953ZZL | ENCFF023NGN |
| ENCFF750EKT | ENCFF895QZG, ENCFF227IZS, ENCFF910IKB, ENCFF937WDE |
| ENCFF140UJB | ENCFF772PJM, ENCFF982BHL |
| ENCFF093QMN | ENCFF304AZH, ENCFF984QXA, ENCFF533FQH, ENCFF836OEO |
| ENCFF439NNR | ENCFF767FSP |
| ENCFF610YXS | ENCFF392XRJ, ENCFF829HZY |
| ENCFF581AZU | ENCFF036ZLP |
| ENCFF474ETY | ENCFF355SGP |
| ENCFF188QGH | ENCFF023NGN |
| ENCFF662HRL | ENCFF306LVM |
| ENCFF416LJL | ENCFF392XRJ, ENCFF829HZY |
| ENCFF958WMW | ENCFF895QZG, ENCFF227IZS, ENCFF910IKB, ENCFF937WDE |
| ENCFF880ZXO | ENCFF510BRO, ENCFF240RBJ |
| ENCFF544ZPF | ENCFF213VVQ, ENCFF063BAN |
| ENCFF544DDY | ENCFF162ZOO, ENCFF968UFH |
| ENCFF623PVZ | ENCFF156FED, ENCFF577FNG |
| ENCFF216TEN | ENCFF299GMC |

| | |
|---|---|
| ENCFF548STV | ENCFF227IZS, ENCFF910IKB |
| ENCFF417FHN | ENCFF156FED, ENCFF577FNG |
| ENCFF035GBL | ENCFF121DUE, ENCFF709XAA |
| ENCFF253HOC | ENCFF547QCM |
| ENCFF568RDF | ENCFF304AZH, ENCFF984QXA, ENCFF533FQH, ENCFF836OEO |
| ENCFF589IXE | ENCFF023NGN |
| ENCFF630JRN | ENCFF023NGN |
| ENCFF691BQZ | ENCFF812TGW, ENCFF234NVU, ENCFF913HVS, ENCFF382XSA |
| ENCFF372NDR | ENCFF308PSW, ENCFF198JCQ |
| ENCFF695TZQ | ENCFF895QZG, ENCFF227IZS, ENCFF910IKB, ENCFF937WDE |
| ENCFF665EXC | ENCFF304AZH, ENCFF984QXA, ENCFF533FQH, ENCFF836OEO |
| ENCFF852HSQ | ENCFF023NGN |
| ENCFF536TPK | ENCFF482LDC |
| ENCFF284MLL | ENCFF604MCA |
| ENCFF834YLT | ENCFF358ZBU |
| ENCFF587HZM | ENCFF895QZG, ENCFF227IZS, ENCFF910IKB, ENCFF937WDE |
| ENCFF986KSB | ENCFF156FED, ENCFF577FNG |
| ENCFF581SOT | ENCFF886VAQ |
| ENCFF353EFN | ENCFF895QZG, ENCFF227IZS, ENCFF910IKB, ENCFF937WDE |
| ENCFF405EME | ENCFF156FED, ENCFF577FNG |
| ENCFF427CAA | ENCFF895QZG, ENCFF227IZS, ENCFF910IKB, ENCFF937WDE |
| ENCFF949ENV | ENCFF002WSA |
| ENCFF378UVV | ENCFF227IZS, ENCFF910IKB |
| ENCFF707JIA | ENCFF227IZS, ENCFF910IKB |
| ENCFF400MMA | ENCFF159FKZ |
| ENCFF554BLC | ENCFF023NGN |
| ENCFF904PBR | ENCFF358ZBU |
| ENCFF183MSQ | ENCFF712WXB, ENCFF790TAN |
| ENCFF236ZUN | ENCFF156FED, ENCFF577FNG |
| ENCFF420DGT | ENCFF712WXB, ENCFF790TAN |
| ENCFF082TOF | ENCFF510BRO, ENCFF240RBJ |
| ENCFF862XFS | ENCFF227IZS, ENCFF910IKB |
| ENCFF215TDO | ENCFF023NGN |
| ENCFF601CVJ | ENCFF227IZS, ENCFF910IKB |
| ENCFF459MNI | ENCFF796JTX, ENCFF720AUK |
| ENCFF491EUN | ENCFF332CUX |
| ENCFF904CPI | ENCFF984QXA, ENCFF836OEO, ENCFF204GLJ, ENCFF304AZH, ENCFF533FQH |

| | |
|---|---|
| ENCFF515YFJ | ENCFF332SVJ |
| ENCFF444ZUH | ENCFF355SGP |
| ENCFF892LAS | ENCFF780MVW |
| ENCFF754GYS | ENCFF895QZG, ENCFF227IZS, ENCFF910IKB, ENCFF937WDE |
| ENCFF564FBJ | ENCFF967YTY |
| ENCFF833ZOK | ENCFF227IZS, ENCFF910IKB |
| ENCFF860JBC | ENCFF392XRJ, ENCFF829HZY |
| ENCFF094FZQ | ENCFF355SGP |
| ENCFF003FNG | ENCFF156FED, ENCFF577FNG |
| ENCFF569DZM | ENCFF772PJM, ENCFF982BHL |
| ENCFF092TPD | ENCFF709XAA, ENCFF332SVJ |
| ENCFF938FQC | ENCFF394JQH |
| ENCFF880SVB | ENCFF227IZS, ENCFF910IKB |
| ENCFF384WCY | ENCFF156FED, ENCFF577FNG |
| ENCFF557QLG | ENCFF824YEY |
| ENCFF827SLL | ENCFF812TGW, ENCFF234NVU, ENCFF913HVS, ENCFF382XSA |
| ENCFF078BWN | ENCFF355SGP |
| ENCFF057KBN | ENCFF156FED, ENCFF577FNG |
| ENCFF587PVQ | ENCFF709XAA, ENCFF332SVJ |
| ENCFF910XWG | ENCFF679QYP |
| ENCFF358SSM | ENCFF895QZG, ENCFF227IZS, ENCFF910IKB, ENCFF937WDE |
| ENCFF170NGB | ENCFF355SGP |
| ENCFF611PXX | ENCFF796JTX, ENCFF720AUK |
| ENCFF079TBV | ENCFF227IZS, ENCFF910IKB |
| ENCFF498EFE | ENCFF390ATQ |
| ENCFF641HZR | ENCFF023NGN |
| ENCFF785NOU | ENCFF712WXB, ENCFF227IZS, ENCFF910IKB, ENCFF790TAN |
| ENCFF135YTU | ENCFF712WXB, ENCFF227IZS, ENCFF910IKB, ENCFF790TAN |
| ENCFF408BKE | ENCFF227IZS, ENCFF910IKB |
| ENCFF715AMR | ENCFF895QZG, ENCFF227IZS, ENCFF910IKB, ENCFF937WDE |
| ENCFF140ZMN | ENCFF023NGN |
| ENCFF639CLN | ENCFF895QZG, ENCFF227IZS, ENCFF910IKB, ENCFF937WDE |
| ENCFF473NMO | ENCFF709XAA, ENCFF332SVJ |
| ENCFF334OKF | ENCFF209VFR |
| ENCFF365BCR | ENCFF023NGN |
| ENCFF391BBO | ENCFF343UNB |
| ENCFF244IJF | ENCFF332CUX |

| | |
|---|---|
| ENCFF285CII | ENCFF441HVC, ENCFF652CFB |
| ENCFF226KOW | ENCFF895QZG, ENCFF227IZS, ENCFF910IKB, ENCFF937WDE |
| ENCFF097GYE | ENCFF023NGN |
| ENCFF362TFV | ENCFF227IZS, ENCFF910IKB |
| ENCFF742TAS | ENCFF709XAA, ENCFF332SVJ |
| ENCFF825VBC | ENCFF448DJP |
| ENCFF239WGU | ENCFF942FFX |
| ENCFF554KEG | ENCFF772PJM, ENCFF982BHL |
| ENCFF318SPV | ENCFF227IZS, ENCFF910IKB |
| ENCFF419SAY | ENCFF355SGP |
| ENCFF492NUF | ENCFF772PJM, ENCFF982BHL |
| ENCFF067TWQ | ENCFF812TGW, ENCFF234NVU, ENCFF913HVS, ENCFF382XSA |
| ENCFF856TBD | ENCFF895QZG, ENCFF227IZS, ENCFF910IKB, ENCFF937WDE |
| ENCFF633CMO | ENCFF355SGP |
| ENCFF983RUX | ENCFF121DUE, ENCFF332SVJ |
| ENCFF550BMQ | ENCFF392XRJ, ENCFF829HZY |
| ENCFF500YYZ | ENCFF895QZG, ENCFF227IZS, ENCFF910IKB, ENCFF937WDE |
| ENCFF562ATM | ENCFF895QZG, ENCFF227IZS, ENCFF910IKB, ENCFF937WDE |
| ENCFF509IQQ | ENCFF392XRJ, ENCFF829HZY |
| ENCFF564KCD | ENCFF709XAA, ENCFF332SVJ |
| ENCFF025QKV | ENCFF772PJM, ENCFF982BHL |
| ENCFF479JUJ | ENCFF332CUX |
| ENCFF221CPW | ENCFF895QZG, ENCFF227IZS, ENCFF910IKB, ENCFF937WDE |
| ENCFF201BQU | ENCFF355SGP |
| ENCFF167NVK | ENCFF895QZG, ENCFF227IZS, ENCFF910IKB, ENCFF937WDE |
| ENCFF157WFG | ENCFF308PSW, ENCFF198JCQ |
| ENCFF124RUL | ENCFF227IZS, ENCFF910IKB |
| ENCFF758LME | ENCFF895QZG, ENCFF227IZS, ENCFF910IKB, ENCFF937WDE |
| ENCFF182YDN | ENCFF895QZG, ENCFF227IZS, ENCFF910IKB, ENCFF937WDE |
| ENCFF161VHM | ENCFF100PTE, ENCFF721LZU |
| ENCFF205JWI | ENCFF895QZG, ENCFF227IZS, ENCFF910IKB, ENCFF937WDE |
| ENCFF490XTX | ENCFF895QZG, ENCFF227IZS, ENCFF910IKB, ENCFF937WDE |
| ENCFF668FCG | ENCFF227IZS, ENCFF910IKB |
| ENCFF500HFO | ENCFF036ZLP |
| ENCFF786GEU | ENCFF276OII |
| ENCFF165STC | ENCFF680XQR |
| ENCFF446HGS | ENCFF712WXB, ENCFF227IZS, ENCFF910IKB, ENCFF790TAN |

| | |
|---|---|
| ENCFF566EXS | ENCFF712WXB, ENCFF790TAN |
| ENCFF954HWS | ENCFF227IZS, ENCFF910IKB |
| ENCFF187ZET | ENCFF121DUE, ENCFF332SVJ |
| ENCFF382DBE | ENCFF332SVJ, ENCFF063BAN |
| ENCFF703CKR | ENCFF895QZG, ENCFF227IZS, ENCFF910IKB, ENCFF937WDE |
| ENCFF923LMN | ENCFF227IZS, ENCFF910IKB |
| ENCFF690YWW | ENCFF568OMP |
| ENCFF895DLK | ENCFF345ODV, ENCFF836GUS |
| ENCFF049RJI | ENCFF227IZS, ENCFF910IKB |
| ENCFF619SRD | ENCFF383YOE |
| ENCFF280AEF | ENCFF002WSA |
| ENCFF497GOO | ENCFF227IZS, ENCFF910IKB |
| ENCFF824KAR | ENCFF584ERB, ENCFF092OLM |
| ENCFF024NZQ | ENCFF895QZG, ENCFF227IZS, ENCFF910IKB, ENCFF937WDE |
| ENCFF532BVM | ENCFF227IZS, ENCFF910IKB |
| ENCFF650RFL | ENCFF240RBJ, ENCFF244GCJ |
| ENCFF060WJH | ENCFF322EZT |
| ENCFF864BUV | ENCFF874TOY |
| ENCFF307UCL | ENCFF712WXB, ENCFF790TAN |
| ENCFF056HVT | ENCFF213VVQ, ENCFF092PMQ |
| ENCFF997BDJ | ENCFF399WOX |
| ENCFF070MZX | ENCFF795UVG |
| ENCFF044TAL | ENCFF772PJM, ENCFF982BHL |
| ENCFF836IMK | ENCFF942FFX |
| ENCFF518PIS | ENCFF772PJM, ENCFF982BHL |
| ENCFF251EVS | ENCFF383YOE |
| ENCFF461OAN | ENCFF635ZZS |
| ENCFF466OLG | ENCFF100PTE, ENCFF721LZU |
| ENCFF709ZMI | ENCFF458FYW |
| ENCFF922JWO | ENCFF243UPF |
| ENCFF178EJY | ENCFF156FED, ENCFF577FNG |
| ENCFF360QAB | ENCFF100PTE, ENCFF721LZU |
| ENCFF179DZP | ENCFF285EWB, ENCFF696ZGZ, ENCFF709XAA |
| ENCFF191EJM | ENCFF115YFR |
| ENCFF350FCE | ENCFF895QZG, ENCFF227IZS, ENCFF910IKB, ENCFF937WDE |
| ENCFF865ISL | ENCFF023NGN |
| ENCFF738QLQ | ENCFF227IZS, ENCFF910IKB |

| | |
|---|---|
| ENCFF413VAR | ENCFF228COG |
| ENCFF225IQS | ENCFF712WXB, ENCFF227IZS, ENCFF910IKB, ENCFF790TAN |
| ENCFF541XEW | ENCFF227IZS, ENCFF910IKB |
| ENCFF635HIS | ENCFF023NGN |
| ENCFF686ZIH | ENCFF895QZG, ENCFF227IZS, ENCFF910IKB, ENCFF937WDE |
| ENCFF562PEW | ENCFF700BKM |
| ENCFF426DLJ | ENCFF355SGP |
| ENCFF540ZTB | ENCFF609HLC |
| ENCFF008CUX | ENCFF712WXB, ENCFF790TAN |
| ENCFF131XTL | ENCFF895QZG, ENCFF227IZS, ENCFF910IKB, ENCFF937WDE |
| ENCFF367OYT | ENCFF345ODV, ENCFF836GUS |
| ENCFF300AUR | ENCFF752BZZ |
| ENCFF620VNP | ENCFF037PYE |
| ENCFF809OVE | ENCFF895QZG, ENCFF227IZS, ENCFF910IKB, ENCFF937WDE |
| ENCFF468LGZ | ENCFF709XAA, ENCFF332SVJ |
| ENCFF398SXO | ENCFF772PJM, ENCFF982BHL |
| ENCFF777ZPZ | ENCFF895QZG, ENCFF227IZS, ENCFF910IKB, ENCFF937WDE |
| ENCFF503IYW | ENCFF156EWJ |
| ENCFF432NAV | ENCFF712WXB, ENCFF790TAN |
| ENCFF284XYU | ENCFF895QZG, ENCFF227IZS, ENCFF910IKB, ENCFF937WDE |
| ENCFF168TBA | ENCFF895QZG, ENCFF227IZS, ENCFF910IKB, ENCFF937WDE |
| ENCFF071KOM | ENCFF156FED, ENCFF577FNG |
| ENCFF655OEX | ENCFF285EWB, ENCFF696ZGZ, ENCFF709XAA, ENCFF204GLJ |
| ENCFF302OZO | ENCFF390ATQ |
| ENCFF204TUQ | ENCFF772PJM, ENCFF982BHL |
| ENCFF339RWU | ENCFF895QZG, ENCFF227IZS, ENCFF910IKB, ENCFF937WDE |
| ENCFF623YPH | ENCFF093MXU |
| ENCFF876GQO | ENCFF332CUX |
| ENCFF561ILM | ENCFF772PJM, ENCFF982BHL |
| ENCFF259RRK | ENCFF332CUX |
| ENCFF334QAB | ENCFF772PJM, ENCFF982BHL |
| ENCFF878UKW | ENCFF247YFL |
| ENCFF433SUG | ENCFF635ZZS |
| ENCFF914ABF | ENCFF942FFX |
| ENCFF406TJQ | ENCFF712WXB, ENCFF227IZS, ENCFF910IKB, ENCFF790TAN |
| ENCFF178YFO | ENCFF895QZG, ENCFF227IZS, ENCFF910IKB, ENCFF937WDE |
| ENCFF102ZOS | ENCFF465FJI |

| | |
|---|---|
| ENCFF047IWR | ENCFF306LVM |
| ENCFF719OIE | ENCFF023NGN |
| ENCFF781JEI | ENCFF392XRJ, ENCFF829HZY |
| ENCFF749LRJ | ENCFF162ZOO, ENCFF092PMQ |
| ENCFF548YYR | ENCFF712WXB, ENCFF790TAN |
| ENCFF016KRJ | ENCFF227IZS, ENCFF910IKB |
| ENCFF625AJO | ENCFF623HUN |
| ENCFF940QAP | ENCFF767FSP |
| ENCFF652HPU | ENCFF392XRJ, ENCFF829HZY |
| ENCFF701KAF | ENCFF243UPF |
| ENCFF057SMG | ENCFF308PSW, ENCFF198JCQ |
| ENCFF207LND | ENCFF227IZS, ENCFF910IKB |
| ENCFF695GTB | ENCFF059OPC |
| ENCFF549MBS | ENCFF712WXB, ENCFF790TAN |
| ENCFF323IVG | ENCFF227IZS, ENCFF910IKB |
| ENCFF698VVG | ENCFF712WXB, ENCFF790TAN |
| ENCFF373YTD | ENCFF712WXB, ENCFF790TAN |
| ENCFF323OVV | ENCFF712WXB, ENCFF790TAN |
| ENCFF022IWG | ENCFF285EWB, ENCFF696ZGZ, ENCFF709XAA |
| ENCFF261BXO | ENCFF121DUE, ENCFF709XAA |
| ENCFF329LWM | ENCFF772PJM, ENCFF982BHL |
| ENCFF338MSY | ENCFF679QYP |
| ENCFF283ZIL | ENCFF285EWB, ENCFF696ZGZ, ENCFF709XAA |
| ENCFF145ZDW | ENCFF073AHU |
| ENCFF550IUC | ENCFF895QZG, ENCFF227IZS, ENCFF910IKB, ENCFF937WDE |
| ENCFF976EMR | ENCFF767FSP |
| ENCFF498KCA | ENCFF266FHE |
| ENCFF340JWK | ENCFF355SGP |
| ENCFF573OTF | ENCFF227IZS, ENCFF910IKB |
| ENCFF917UQJ | ENCFF355SGP |
| ENCFF079SPF | ENCFF227IZS, ENCFF910IKB |
| ENCFF921KTI | ENCFF355SGP |
| ENCFF618MHM | ENCFF936VXD, ENCFF092OEQ |
| ENCFF253JTJ | ENCFF895QZG, ENCFF227IZS, ENCFF910IKB, ENCFF937WDE |
| ENCFF202BLM | ENCFF213VVQ, ENCFF063BAN |
| ENCFF931HHQ | ENCFF895QZG, ENCFF227IZS, ENCFF910IKB, ENCFF937WDE |
| ENCFF651HPM | ENCFF772PJM, ENCFF982BHL |

| | |
|---|---|
| ENCFF331TRC | ENCFF812TGW, ENCFF234NVU, ENCFF913HVS, ENCFF382XSA |
| ENCFF855NIZ | ENCFF895QZG, ENCFF227IZS, ENCFF910IKB, ENCFF937WDE |
| ENCFF122TGF | ENCFF227IZS, ENCFF910IKB |
| ENCFF700WSI | ENCFF895QZG, ENCFF227IZS, ENCFF910IKB, ENCFF937WDE |
| ENCFF050LIC | ENCFF482LDC |
| ENCFF360QBV | ENCFF023NGN |
| ENCFF400SKV | ENCFF266FHE |
| ENCFF643UCP | ENCFF332CUX |
| ENCFF323MTY | ENCFF767FSP |
| ENCFF546IMN | ENCFF772PJM, ENCFF982BHL |
| ENCFF553WMU | ENCFF482LDC |
| ENCFF576EDK | ENCFF156FED, ENCFF577FNG |
| ENCFF821HMU | ENCFF227IZS, ENCFF910IKB |
| ENCFF488BQX | ENCFF227IZS, ENCFF910IKB |
| ENCFF129RYE | ENCFF712WXB, ENCFF790TAN |
| ENCFF693JVP | ENCFF037PYE |
| ENCFF068XVC | ENCFF023NGN |
| ENCFF525CPA | ENCFF895QZG, ENCFF227IZS, ENCFF910IKB, ENCFF937WDE |
| ENCFF540VVE | ENCFF156EWJ |
| ENCFF400BSN | ENCFF023NGN |
| ENCFF726AYE | ENCFF156FED, ENCFF577FNG |
| ENCFF403WNR | ENCFF767FSP |
| ENCFF040MZK | ENCFF023NGN |
| ENCFF642RHD | ENCFF355SGP |
| ENCFF289RGM | ENCFF304AZH, ENCFF984QXA, ENCFF533FQH, ENCFF836OEO |
| ENCFF512YJL | ENCFF162ZOO, ENCFF332SVJ |
| ENCFF337GMY | ENCFF227IZS, ENCFF910IKB |
| ENCFF786KLU | ENCFF596JNR |
| ENCFF180ZQP | ENCFF568OMP |
| ENCFF778PDI | ENCFF772PJM, ENCFF982BHL |
| ENCFF677JTW | ENCFF895QZG, ENCFF227IZS, ENCFF910IKB, ENCFF937WDE |
| ENCFF788ICY | ENCFF355SGP |
| ENCFF247FGI | ENCFF227IZS, ENCFF910IKB |
| ENCFF774RUP | ENCFF780MVW |
| ENCFF198RTF | ENCFF121DUE, ENCFF332SVJ |
| ENCFF431WVX | ENCFF712WXB, ENCFF227IZS, ENCFF910IKB, ENCFF790TAN |
| ENCFF784SCN | ENCFF772PJM, ENCFF982BHL |

| | |
|---|---|
| ENCFF662KUU | ENCFF898KHD |
| ENCFF327XSI | ENCFF394JQH |
| ENCFF066JMV | ENCFF712WXB, ENCFF790TAN |
| ENCFF423MBY | ENCFF712WXB, ENCFF790TAN |
| ENCFF151GOI | ENCFF023NGN |
| ENCFF602UAC | ENCFF240RBJ, ENCFF244GCJ |
| ENCFF103HTZ | ENCFF874TOY |
| ENCFF002PNZ | ENCFF779PRW |
| ENCFF149LTH | ENCFF895QZG, ENCFF227IZS, ENCFF910IKB, ENCFF937WDE |
| ENCFF975ZXB | ENCFF023NGN |
| ENCFF299HHL | ENCFF942FFX |
| ENCFF548NQU | ENCFF942FFX |
| ENCFF465ZZG | ENCFF604MCA |
| ENCFF303ICH | ENCFF712WXB, ENCFF227IZS, ENCFF910IKB, ENCFF790TAN |
| ENCFF773DBT | ENCFF547QCM |
| ENCFF729WGC | ENCFF023NGN |
| ENCFF982AFE | ENCFF023NGN |
| ENCFF397LNB | ENCFF895QZG, ENCFF227IZS, ENCFF910IKB, ENCFF937WDE |
| ENCFF883MZR | ENCFF895QZG, ENCFF227IZS, ENCFF910IKB, ENCFF937WDE |
| ENCFF417HNM | ENCFF156FED, ENCFF577FNG |
| ENCFF125HKL | ENCFF227IZS, ENCFF910IKB |
| ENCFF396AXY | ENCFF343UNB |
| ENCFF680KUB | ENCFF756KAA |
| ENCFF213JOK | ENCFF023NGN |
| ENCFF198CVB | ENCFF023NGN |
| ENCFF502VUK | ENCFF285EWB, ENCFF696ZGZ, ENCFF709XAA, ENCFF204GLJ |
| ENCFF378QWS | ENCFF247YFL |
| ENCFF803TYM | ENCFF712WXB, ENCFF790TAN |
| ENCFF976DWY | ENCFF023NGN |
| ENCFF589UVV | ENCFF322EZT |
| ENCFF408GAN | ENCFF661ZRO, ENCFF872FDX |
| ENCFF799FNR | ENCFF895QZG, ENCFF227IZS, ENCFF910IKB, ENCFF937WDE |
| ENCFF171VBC | ENCFF023NGN |
| ENCFF477CYZ | ENCFF895QZG, ENCFF227IZS, ENCFF910IKB, ENCFF937WDE |
| ENCFF564CXM | ENCFF227IZS, ENCFF910IKB |
| ENCFF725WJS | ENCFF227IZS, ENCFF910IKB |
| ENCFF677BBS | ENCFF121DUE, ENCFF332SVJ |

| | |
|---|---|
| ENCFF294DBF | ENCFF209VFR |
| ENCFF784MLU | ENCFF942FFX |
| ENCFF308RGN | ENCFF772PJM, ENCFF982BHL |
| ENCFF019RWB | ENCFF895QZG, ENCFF227IZS, ENCFF910IKB, ENCFF937WDE |
| ENCFF019PLG | ENCFF156FED, ENCFF577FNG |
| ENCFF555QMG | ENCFF227IZS, ENCFF910IKB |
| ENCFF990RGE | ENCFF895QZG, ENCFF227IZS, ENCFF910IKB, ENCFF937WDE |
| ENCFF502NAY | ENCFF895QZG, ENCFF227IZS, ENCFF910IKB, ENCFF937WDE |
| ENCFF126SXE | ENCFF772PJM, ENCFF982BHL |
| ENCFF593XTM | ENCFF159FKZ |
| ENCFF846CYU | ENCFF895QZG, ENCFF227IZS, ENCFF910IKB, ENCFF937WDE |
| ENCFF156NIH | ENCFF355SGP |
| ENCFF856EXS | ENCFF699KKU |
| ENCFF494VZW | ENCFF873PSH |
| ENCFF614RBF | ENCFF156FED, ENCFF577FNG |
| ENCFF850FRG | ENCFF772PJM, ENCFF982BHL |
| ENCFF606LUD | ENCFF227IZS, ENCFF910IKB |
| ENCFF682NGE | ENCFF100PTE, ENCFF721LZU |
| ENCFF730MAX | ENCFF647OZY |
| ENCFF358HOW | ENCFF712WXB, ENCFF790TAN |
| ENCFF145NON | ENCFF227IZS, ENCFF910IKB |
| ENCFF046VNU | ENCFF227IZS, ENCFF910IKB |
| ENCFF719XQY | ENCFF162ZOO, ENCFF968UFH |
| ENCFF437UWB | ENCFF895QZG, ENCFF227IZS, ENCFF910IKB, ENCFF937WDE |
| ENCFF715AOC | ENCFF227IZS, ENCFF910IKB |
| ENCFF158BYF | ENCFF824YEY |
| ENCFF339YBR | ENCFF772PJM, ENCFF982BHL |
| ENCFF606TJM | ENCFF230BZG |
| ENCFF401PIJ | ENCFF308PSW, ENCFF198JCQ |
| ENCFF563DGZ | ENCFF895QZG, ENCFF227IZS, ENCFF910IKB, ENCFF937WDE |
| ENCFF304KSV | ENCFF712WXB, ENCFF790TAN |
| ENCFF540DXB | ENCFF023NGN |
| ENCFF033UID | ENCFF332SVJ, ENCFF505ZUJ |
| ENCFF205VDU | ENCFF895QZG, ENCFF227IZS, ENCFF910IKB, ENCFF937WDE |
| ENCFF811CBJ | ENCFF228COG |
| ENCFF871KNP | ENCFF227IZS, ENCFF910IKB |
| ENCFF450TNG | ENCFF227IZS, ENCFF910IKB |

| | |
|---|---|
| ENCFF615CWW | ENCFF767FSP |
| ENCFF810IRF | ENCFF423TQJ |
| ENCFF056XVW | ENCFF162ZOO, ENCFF092PMQ |
| ENCFF871GXE | ENCFF392XRJ, ENCFF829HZY |
| ENCFF694KHP | ENCFF680XQR |
| ENCFF203MUO | ENCFF936VXD, ENCFF092OEQ |
| ENCFF321ZQU | ENCFF023NGN |
| ENCFF294UTM | ENCFF895QZG, ENCFF227IZS, ENCFF910IKB, ENCFF937WDE |
| ENCFF888AOR | ENCFF332SVJ, ENCFF063BAN |
| ENCFF706XFF | ENCFF796JTX, ENCFF720AUK |
| ENCFF649ORS | ENCFF868RUE |
| ENCFF387BDW | ENCFF895QZG, ENCFF227IZS, ENCFF910IKB, ENCFF937WDE |
| ENCFF213CZL | ENCFF392XRJ, ENCFF829HZY |
| ENCFF728WOA | ENCFF482LDC |
| ENCFF072SKI | ENCFF332SVJ |
| ENCFF434YXL | ENCFF868RUE |
| ENCFF680COV | ENCFF772PJM, ENCFF982BHL |
| ENCFF154YTN | ENCFF304AZH, ENCFF984QXA, ENCFF533FQH, ENCFF836OEO |
| ENCFF263MBV | ENCFF895QZG, ENCFF227IZS, ENCFF910IKB, ENCFF937WDE |
| ENCFF597WPX | ENCFF699KKU |
| ENCFF412DRO | ENCFF332SVJ, ENCFF505ZUJ |
| ENCFF634KQT | ENCFF647SZO |
| ENCFF191CWT | ENCFF482LDC |
| ENCFF089XMI | ENCFF332CUX |
| ENCFF659KOL | ENCFF683DQU, ENCFF240RBJ |
| ENCFF980YNA | ENCFF967YTY |
| ENCFF829XLC | ENCFF936BLQ |
| ENCFF577WGL | ENCFF177OCL |
| ENCFF454MMY | ENCFF772PJM, ENCFF982BHL |
| ENCFF011OSI | ENCFF355SGP |
| ENCFF831ODM | ENCFF073AHU |
| ENCFF139WUV | ENCFF355SGP |
| ENCFF013MXK | ENCFF895QZG, ENCFF227IZS, ENCFF910IKB, ENCFF937WDE |
| ENCFF794ABP | ENCFF772PJM, ENCFF982BHL |
| ENCFF874YYW | ENCFF355SGP |
| ENCFF613CDR | ENCFF355SGP |
| ENCFF066TDG | ENCFF162ZOO, ENCFF332SVJ |

| | |
|---|---|
| ENCFF200PYZ | ENCFF023NGN |
| ENCFF158ZVB | ENCFF767FSP |
| ENCFF799RPS | ENCFF156FED, ENCFF577FNG |
| ENCFF115BME | ENCFF059OPC |
| ENCFF795OAZ | ENCFF895QZG, ENCFF227IZS, ENCFF910IKB, ENCFF937WDE |
| ENCFF353XMB | ENCFF895QZG, ENCFF227IZS, ENCFF910IKB, ENCFF937WDE |
| ENCFF081HVQ | ENCFF873PSH |
| ENCFF168KTM | ENCFF942FFX |
| ENCFF945TUI | ENCFF886VAQ |
| ENCFF790FNF | ENCFF304AZH, ENCFF984QXA, ENCFF533FQH, ENCFF836OEO |
| ENCFF771KEQ | ENCFF936BLQ |
| ENCFF355PQK | ENCFF023NGN |
| ENCFF030XLK | ENCFF332SVJ |
| ENCFF925CYF | ENCFF156FED, ENCFF577FNG |
| ENCFF873BRB | ENCFF895QZG, ENCFF227IZS, ENCFF910IKB, ENCFF937WDE |
| ENCFF837ZOY | ENCFF712WXB, ENCFF790TAN |
| ENCFF784MTZ | ENCFF895QZG, ENCFF227IZS, ENCFF910IKB, ENCFF937WDE |
| ENCFF979BFW | ENCFF465FJI |
| ENCFF010TVY | ENCFF308PSW, ENCFF198JCQ |
| ENCFF483RLD | ENCFF767FSP |
| ENCFF425MJK | ENCFF156FED, ENCFF577FNG |
| ENCFF221XVM | ENCFF274BOZ |
| ENCFF393ZAK | ENCFF772PJM, ENCFF982BHL |
| ENCFF891GFG | ENCFF355SGP |
| ENCFF599IYT | ENCFF712WXB, ENCFF790TAN |
| ENCFF586KRI | ENCFF898KHD |
| ENCFF218JFL | ENCFF767FSP |
| ENCFF385PRZ | ENCFF156FED, ENCFF577FNG |
| ENCFF954PRL | ENCFF392XRJ, ENCFF829HZY |
| ENCFF149WGG | ENCFF712WXB, ENCFF790TAN |
| ENCFF226IAA | ENCFF895QZG, ENCFF227IZS, ENCFF910IKB, ENCFF937WDE |
| ENCFF090SFR | ENCFF162ZOO, ENCFF332SVJ |
| ENCFF257QQX | ENCFF156FED, ENCFF577FNG |
| ENCFF786TLS | ENCFF683DQU, ENCFF240RBJ |
| ENCFF330XAC | ENCFF712WXB, ENCFF790TAN |
| ENCFF780DBH | ENCFF332SVJ |
| ENCFF077NYA | ENCFF712WXB, ENCFF790TAN |

| | |
|---|---|
| ENCFF793KZB | ENCFF984QXA, ENCFF836OEO, ENCFF204GLJ, ENCFF304AZH, ENCFF533FQH |
| ENCFF006GQZ | ENCFF023NGN |
| ENCFF471TDC | ENCFF100PTE, ENCFF721LZU |
| ENCFF563FZR | ENCFF895QZG, ENCFF227IZS, ENCFF910IKB, ENCFF937WDE |
| ENCFF857YYV | ENCFF895QZG, ENCFF227IZS, ENCFF910IKB, ENCFF937WDE |
| ENCFF988VSQ | ENCFF023NGN |
| ENCFF722LJA | ENCFF227IZS, ENCFF910IKB |
| ENCFF525JGC | ENCFF100PTE, ENCFF721LZU |
| ENCFF670SFK | ENCFF102ZZG |
| ENCFF786MHJ | ENCFF895QZG, ENCFF227IZS, ENCFF910IKB, ENCFF937WDE |
| ENCFF350NUJ | ENCFF227IZS, ENCFF910IKB |
| ENCFF772XIR | ENCFF227IZS, ENCFF910IKB |
| ENCFF201MXQ | ENCFF227IZS, ENCFF910IKB |
| ENCFF616QUW | ENCFF177OCL |
| ENCFF836NST | ENCFF332SVJ, ENCFF204GLJ |
| ENCFF924CYX | ENCFF023NGN |
| ENCFF507UJN | ENCFF162ZOO, ENCFF092PMQ |
| ENCFF559BJS | ENCFF227IZS, ENCFF910IKB |
| ENCFF697TDR | ENCFF172LMI |
| ENCFF035CJB | ENCFF895QZG, ENCFF227IZS, ENCFF910IKB, ENCFF937WDE |
| ENCFF823GCX | ENCFF812TGW, ENCFF234NVU, ENCFF913HVS, ENCFF382XSA |
| ENCFF728IYF | ENCFF355SGP |
| ENCFF636IGK | ENCFF895QZG, ENCFF227IZS, ENCFF910IKB, ENCFF937WDE |
| ENCFF760JNE | ENCFF423TQJ |
| ENCFF007ALP | ENCFF227IZS, ENCFF910IKB |
| ENCFF302WSZ | ENCFF392XRJ, ENCFF829HZY |
| ENCFF723QZO | ENCFF156FED, ENCFF577FNG |
| ENCFF995UTD | ENCFF227IZS, ENCFF910IKB |
| ENCFF361MTW | ENCFF100PTE, ENCFF721LZU |
| ENCFF027QPB | ENCFF172LMI |
| ENCFF535SSL | ENCFF816BIC |
| ENCFF904OXK | ENCFF712WXB, ENCFF790TAN |
| ENCFF108SDD | ENCFF895QZG, ENCFF227IZS, ENCFF910IKB, ENCFF937WDE |
| ENCFF036IOO | ENCFF712WXB, ENCFF790TAN |
| ENCFF490ICL | ENCFF227IZS, ENCFF910IKB |
| ENCFF011XRF | ENCFF285EWB, ENCFF696ZGZ, ENCFF709XAA |
| ENCFF401BQM | ENCFF772PJM, ENCFF982BHL |

| | |
|---|---|
| ENCFF275XJW | ENCFF308PSW, ENCFF198JCQ |
| ENCFF931HWG | ENCFF700BKM |
| ENCFF900QPQ | ENCFF213VVQ, ENCFF092PMQ |
| ENCFF702WHX | ENCFF895QZG, ENCFF227IZS, ENCFF910IKB, ENCFF937WDE |
| ENCFF257OOQ | ENCFF712WXB, ENCFF227IZS, ENCFF910IKB, ENCFF790TAN |
| ENCFF918CSI | ENCFF156FED, ENCFF577FNG |
| ENCFF041ZVD | ENCFF712WXB, ENCFF227IZS, ENCFF910IKB, ENCFF790TAN |
| ENCFF717OYW | ENCFF895QZG, ENCFF227IZS, ENCFF910IKB, ENCFF937WDE |
| ENCFF314OQP | ENCFF304AZH, ENCFF984QXA, ENCFF533FQH, ENCFF836OEO |
| ENCFF903JXA | ENCFF285EWB, ENCFF696ZGZ, ENCFF709XAA |
| ENCFF495MEU | ENCFF023NGN |
| ENCFF703YNU | ENCFF767FSP |
| ENCFF802CIM | ENCFF227IZS, ENCFF910IKB |
| ENCFF212PQL | ENCFF623HUN |
| ENCFF880ODJ | ENCFF156FED, ENCFF577FNG |
| ENCFF342FIU | ENCFF712WXB, ENCFF790TAN |
| ENCFF489YJG | ENCFF482LDC |
| ENCFF726UVN | ENCFF023NGN |
| ENCFF567GMS | ENCFF812TGW, ENCFF234NVU, ENCFF913HVS, ENCFF382XSA |
| ENCFF516UTV | ENCFF304AZH, ENCFF984QXA, ENCFF533FQH, ENCFF836OEO |
| ENCFF738AVH | ENCFF895QZG, ENCFF227IZS, ENCFF910IKB, ENCFF937WDE |
| ENCFF438XEX | ENCFF458FYW |
| ENCFF771NSF | ENCFF355SGP |
| ENCFF267EEY | ENCFF227IZS, ENCFF910IKB |
| ENCFF584HKJ | ENCFF772PJM, ENCFF982BHL |
| ENCFF106BHY | ENCFF093MXU |
| ENCFF757TIT | ENCFF756KAA |
| ENCFF072KEJ | ENCFF227IZS, ENCFF910IKB |
| ENCFF535PLW | ENCFF812TGW, ENCFF234NVU, ENCFF913HVS, ENCFF382XSA |
| ENCFF797FVU | ENCFF213VVQ, ENCFF063BAN |
| ENCFF530UEA | ENCFF895QZG, ENCFF227IZS, ENCFF910IKB, ENCFF937WDE |
| ENCFF315DCZ | ENCFF712WXB, ENCFF790TAN |
| ENCFF494OJC | ENCFF895QZG, ENCFF227IZS, ENCFF910IKB, ENCFF937WDE |
| ENCFF227TEG | ENCFF321YGO |
| ENCFF346SNT | ENCFF779PRW |
| ENCFF207NLX | ENCFF709XAA, ENCFF332SVJ |
| ENCFF814CHV | ENCFF482LDC |

| | |
|---|---|
| ENCFF373AFZ | ENCFF895QZG, ENCFF227IZS, ENCFF910IKB, ENCFF937WDE |
| ENCFF752FIR | ENCFF895QZG, ENCFF227IZS, ENCFF910IKB, ENCFF937WDE |
| ENCFF391HFU | ENCFF812TGW, ENCFF234NVU, ENCFF913HVS, ENCFF382XSA |
| ENCFF195WOK | ENCFF895QZG, ENCFF227IZS, ENCFF910IKB, ENCFF937WDE |
| ENCFF632WSK | ENCFF895QZG, ENCFF227IZS, ENCFF910IKB, ENCFF937WDE |
| ENCFF587WWS | ENCFF772PJM, ENCFF982BHL |
| ENCFF388QNA | ENCFF023NGN |
| ENCFF156OFZ | ENCFF100PTE, ENCFF721LZU |
| ENCFF031YZL | ENCFF100PTE, ENCFF721LZU |
| ENCFF357MHM | ENCFF332CUX |
| ENCFF919HAZ | ENCFF772PJM, ENCFF982BHL |
| ENCFF229VMU | ENCFF227IZS, ENCFF910IKB |
| ENCFF860IRS | ENCFF895QZG, ENCFF227IZS, ENCFF910IKB, ENCFF937WDE |
| ENCFF914TEL | ENCFF227IZS, ENCFF910IKB |
| ENCFF420ZVF | ENCFF023NGN |
| ENCFF050VHP | ENCFF712WXB, ENCFF790TAN |
| ENCFF331BZT | ENCFF772PJM, ENCFF982BHL |
| ENCFF908PSF | ENCFF276OII |
| ENCFF546ZLD | ENCFF895QZG, ENCFF227IZS, ENCFF910IKB, ENCFF937WDE |
| ENCFF397ZVS | ENCFF984QXA, ENCFF836OEO, ENCFF204GLJ, ENCFF304AZH, ENCFF533FQH |
| ENCFF769DDY | ENCFF895QZG, ENCFF227IZS, ENCFF910IKB, ENCFF937WDE |
| ENCFF401KIO | ENCFF772PJM, ENCFF982BHL |
| ENCFF657TZQ | ENCFF895QZG, ENCFF227IZS, ENCFF910IKB, ENCFF937WDE |
| ENCFF014DRH | ENCFF482LDC |
| ENCFF157HYJ | ENCFF227IZS, ENCFF910IKB |
| ENCFF364SJK | ENCFF796JTX, ENCFF720AUK |
| ENCFF320WXN | ENCFF895QZG, ENCFF227IZS, ENCFF910IKB, ENCFF937WDE |
| ENCFF273NVY | ENCFF227IZS, ENCFF910IKB |
| ENCFF540SXT | ENCFF162ZOO, ENCFF092PMQ |
| ENCFF268CBK | ENCFF895QZG, ENCFF227IZS, ENCFF910IKB, ENCFF937WDE |
| ENCFF503JQK | ENCFF100PTE, ENCFF721LZU |
| ENCFF795XBH | ENCFF795UVG |
| ENCFF687QLR | ENCFF115YFR |
| ENCFF438GBD | ENCFF227IZS, ENCFF910IKB |
| ENCFF487SUP | ENCFF895QZG, ENCFF227IZS, ENCFF910IKB, ENCFF937WDE |
| ENCFF641OBF | ENCFF661ZRO, ENCFF872FDX |
| ENCFF488CXC | ENCFF023NGN |

| | |
|---|---|
| ENCFF008IZS | ENCFF712WXB, ENCFF790TAN |
| ENCFF961HAZ | ENCFF392XRJ, ENCFF829HZY |
| ENCFF594OID | ENCFF023NGN |
| ENCFF656BKZ | ENCFF102ZZG |
| ENCFF779IQL | ENCFF712WXB, ENCFF227IZS, ENCFF910IKB, ENCFF790TAN |
| ENCFF908MXL | ENCFF162ZOO, ENCFF092PMQ |
| ENCFF951ITB | ENCFF332CUX |
| ENCFF762TAN | ENCFF308PSW, ENCFF198JCQ |
| ENCFF749RRI | ENCFF942FFX |
| ENCFF964COC | ENCFF023NGN |
| ENCFF384DMV | ENCFF647OZY |
| ENCFF254IUH | ENCFF895QZG, ENCFF227IZS, ENCFF910IKB, ENCFF937WDE |
| ENCFF477DKP | ENCFF023NGN |
| ENCFF744TCZ | ENCFF895QZG, ENCFF227IZS, ENCFF910IKB, ENCFF937WDE |
| ENCFF779CRV | ENCFF355SGP |
| ENCFF014UUB | ENCFF023NGN |
| ENCFF928JGK | ENCFF895QZG, ENCFF227IZS, ENCFF910IKB, ENCFF937WDE |
| ENCFF456VOS | ENCFF392XRJ, ENCFF829HZY |
| ENCFF428XVO | ENCFF392XRJ, ENCFF829HZY |
| ENCFF315DZX | ENCFF156FED, ENCFF577FNG |
| ENCFF946HWT | ENCFF023NGN |
| ENCFF285FGV | ENCFF399WOX |
| ENCFF042LKP | ENCFF712WXB, ENCFF790TAN |
| ENCFF527CTN | ENCFF767FSP |
| ENCFF463LMJ | ENCFF767FSP |
| ENCFF015UIX | ENCFF895QZG, ENCFF227IZS, ENCFF910IKB, ENCFF937WDE |
| ENCFF933HFV | ENCFF895QZG, ENCFF227IZS, ENCFF910IKB, ENCFF937WDE |
| ENCFF434RFX | ENCFF023NGN |
| ENCFF448SJG | ENCFF596JNR |
| ENCFF046ACV | ENCFF895QZG, ENCFF227IZS, ENCFF910IKB, ENCFF937WDE |
| ENCFF096KBP | ENCFF441HVC, ENCFF652CFB |
| ENCFF812KIP | ENCFF285EWB, ENCFF696ZGZ, ENCFF709XAA |
| ENCFF033JFM | ENCFF227IZS, ENCFF910IKB |

Table 3: Table for the transcription factors (TFs) and their corresponding motif ID from JASPAR.

| TF | ID |
|---|---|
| MXI1 | MA1108.1 |
| THAP1 | MA0597.1 |
| TFDP1 | MA1122.1 |
| Gata1 | MA0035.2 |
| ZNF384 | MA1125.1 |
| Tcf12 | MA0521.1 |
| MITF | MA0620.2 |
| E2F4 | MA0470.1 |
| MAFF | MA0495.1 |
| CEBPB | MA0466.1 |
| NR2F2 | MA1111.1 |
| TEAD2 | MA1121.1 |
| JUNB | MA0490.1 |
| CTCFL | MA1102.1 |
| YY1 | MA0095.2 |
| USF1 | MA0093.2 |
| REST | MA0138.2 |
| USF2 | MA0526.1 |
| Rfx1 | MA0509.1 |
| Zfx | MA0146.1 |
| ESRRA | MA0592.1 |
| Myc | MA0147.1 |
| ELF1 | MA0473.1 |
| STAT1 | MA0137.2 |
| ZNF24 | MA1124.1 |
| ZBTB7A | MA0750.2 |
| IRF1 | MA0050.2 |
| JUN | MA0488.1 |
| GATA2 | MA0036.2 |
| TCF7L2 | MA0523.1 |
| SP1 | MA0079.3 |
| CTCF | MA0139.1 |
| NR4A1 | MA1112.1 |
| NR2C2 | MA0504.1 |

| | |
|---|---|
| NRF1 | MA0506.1 |
| FOXA1 | MA0148.1 |
| FOXK2 | MA1103.1 |
| JUND | MA0491.1 |
| E2F1 | MA0024.2 |
| E2F6 | MA0471.1 |
| ZNF263 | MA0528.1 |
| ZBTB33 | MA0527.1 |
| Ets1 | MA0098.2 |
| Gabpa | MA0062.2 |
| RUNX1 | MA0002.2 |
| MAX | MA0058.2 |
| FOSL1 | MA0477.1 |

Table 4: Table for the ChIP-seq experiments and their corresponding ChIP-seq replicate samples, TFs and controls for the A549 cell line from the ENCODE database used in our analysis.

| Experiment | Replicates | TF | Controls |
|---|---|---|---|
| ENCSR182OZC | ENCFF791DRP<br>ENCFF217RBI | CEBPB | ENCFF634ULC<br>ENCFF632UPH<br>ENCFF368OTV |
| ENCSR375BUB | ENCFF073MBT<br>ENCFF280ZFT | CEBPB | ENCFF214UMU<br>ENCFF773DUX<br>ENCFF408NFU |
| ENCSR606ZTC | ENCFF347MNU<br>ENCFF417GPF | CEBPB | ENCFF455UAB<br>ENCFF887YTT<br>ENCFF081TBO |
| ENCSR623KNM | ENCFF757GXN<br>ENCFF826GGN | ELK1 | ENCFF949XNJ<br>ENCFF918AJW |
| ENCSR000BQO | ENCFF125MJO<br>ENCFF585INN | FOSL2 | ENCFF656HEF |
| ENCSR419TWL | ENCFF504YVD<br>ENCFF595EIS | HES2 | ENCFF634ULC<br>ENCFF632UPH<br>ENCFF368OTV |
| ENCSR991VVW | ENCFF476XBN<br>ENCFF761UEZ | JUN | ENCFF193ABY<br>ENCFF222ACA<br>ENCFF639UDD |
| ENCSR269RPR | ENCFF179XAQ<br>ENCFF330XFU | JUNB | ENCFF171YYX<br>ENCFF631DES<br>ENCFF298EPS |
| ENCSR431LRW | ENCFF599JTK<br>ENCFF389OFH | JUNB | ENCFF653HKQ<br>ENCFF097CSC<br>ENCFF987XCE |
| ENCSR892DRK | ENCFF364NWO<br>ENCFF808ADX | REST | ENCFF572IKT<br>ENCFF714AMB |

Table 5: Table for the ChIP-seq experiments and their corresponding ChIP-seq replicate samples, TFs and controls for the GM12878 cell line from the ENCODE database used in our analysis.

| Experiment | Replicates | TF | Controls |
|---|---|---|---|
| ENCSR841NDX | ENCFF028TNY<br>ENCFF444PPF | ELF1 | ENCFF666ATR<br>ENCFF322NTO |
| ENCSR000BMB | ENCFF739SRY<br>ENCFF735DGJ | ELF1 | ENCFF488YYE |
| ENCSR000DZB | ENCFF211VKF<br>ENCFF784XUE | ELK1 | ENCFF477ZKJ<br>ENCFF450WED<br>ENCFF824NQO<br>ENCFF710SMS<br>ENCFF579QDW<br>ENCFF813LMQ |
| ENCSR000BGY | ENCFF240MQI<br>ENCFF888PAI | IRF4 | ENCFF562HPN<br>ENCFF100EIH<br>ENCFF438FFV |
| ENCSR000BQL | ENCFF983YCI<br>ENCFF207QTV | NFATC1 | ENCFF754WTG<br>ENCFF966AVZ<br>ENCFF537DAJ |
| ENCSR000BGR | ENCFF791EPM<br>ENCFF845MYC | PBX3 | ENCFF562HPN<br>ENCFF100EIH<br>ENCFF438FFV |
| ENCSR000BQS | ENCFF894EID<br>ENCFF569QEN | REST | ENCFF430ZCF<br>ENCFF100EIH<br>ENCFF438FFV<br>ENCFF562HPN |
| ENCSR000BRI | ENCFF884LEJ<br>ENCFF579PRC | RUNX3 | ENCFF754WTG<br>ENCFF966AVZ<br>ENCFF537DAJ |
| ENCSR000BGE | ENCFF263NOT<br>ENCFF731ZNW | SRF | ENCFF862QZT<br>ENCFF289ONG |
| ENCSR000BGI | ENCFF737VAT<br>ENCFF074OYP | USF | ENCFF862QZT<br>ENCFF289ONG |

Table 6: Table for the ChIP-seq experiments and their corresponding ChIP-seq replicate samples, TFs and controls for the HepG2 cell line from the ENCODE database used in our analysis.

| Experiment | Replicates | TF | Controls |
|---|---|---|---|
| ENCSR000BQI | ENCFF090KCF<br>ENCFF499BWX | CEBPB | ENCFF175NMQ<br>ENCFF285LVE |
| ENCSR000EEE | ENCFF677PSB<br>ENCFF514FUP | CEBPB | ENCFF165KZY |
| ENCSR000BIE | ENCFF435DKZ<br>ENCFF178SXE | CTCF | ENCFF175NMQ<br>ENCFF285LVE |
| ENCSR112ALD | ENCFF011HOS<br>ENCFF320SCI | CREB1 | ENCFF950AXC<br>ENCFF190EPQ |
| ENCSR267DFA | ENCFF396NXZ<br>ENCFF988UCQ | FOXA1 | ENCFF950AXC<br>ENCFF190EPQ |
| ENCSR000BMO | ENCFF332SRJ<br>ENCFF401YVR | FOXA1 | ENCFF175NMQ<br>ENCFF193YIO<br>ENCFF285LVE<br>ENCFF943DZB |
| ENCSR000BHP | ENCFF953PCA<br>ENCFF185DVY | FOSL2 | ENCFF943DZB<br>ENCFF193YIO |
| ENCSR000BMZ | ENCFF930EXY<br>ENCFF418EVV | ELF1 | ENCFF175NMQ<br>ENCFF285LVE |
| ENCSR000BJL | ENCFF195HUS<br>ENCFF291BBG | REST | ENCFF249PQD<br>ENCFF741TQN |
| ENCSR000EEK | ENCFF074GYD<br>ENCFF122BQB | JUN | ENCFF165KZY |

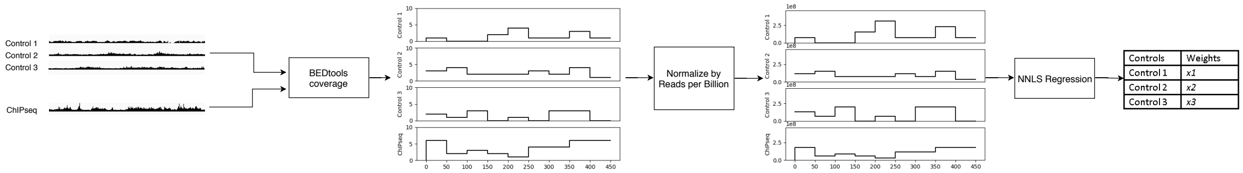


Figure 1: Flowchart for the estimation of weights per control.

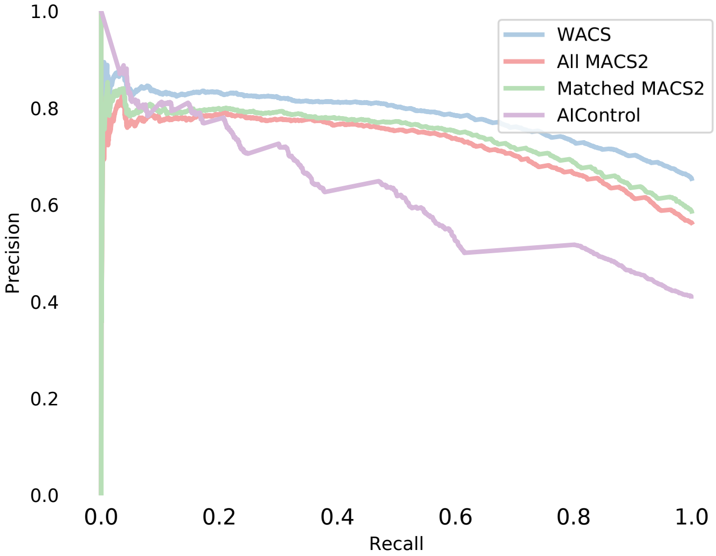


Figure 2: Example of precision recall curve for TF ZNF24 ChIP-seq dataset ENCFF109OWW.

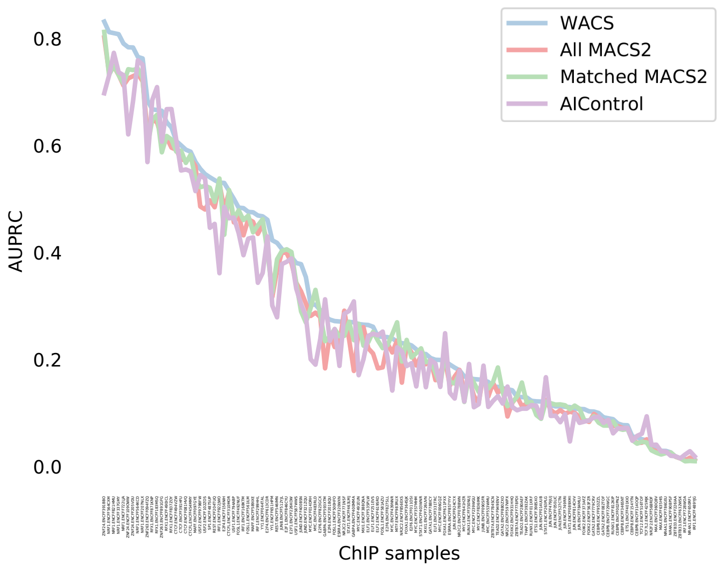


Figure 3: AUPRC for the treatment samples.

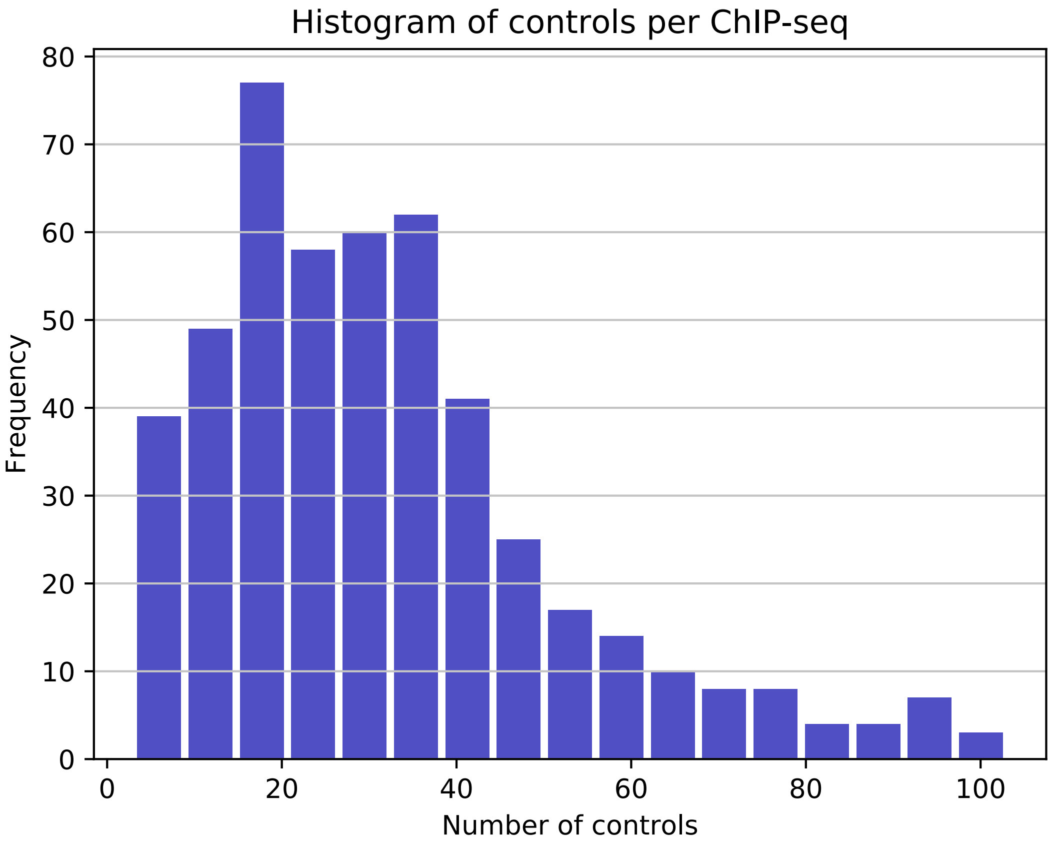


Figure 4: Histogram of the overall number of control used per ChIP-seq dataset using WACS.
